# Knockout of P2Y12 receptor facilitates microglia-neuron body-to-body interactions and accelerates prion disease

**DOI:** 10.1101/2025.04.07.647619

**Authors:** Natallia Makarava, Tarek Safadi, Olga Bocharova, Olga Mychko, Narayan P. Pandit, Kara Molesworth, Ukpong B. Eyo, Ilia V. Baskakov

**Author notes:** Correspondence: 111 Penn St., Baltimore, MD, 21201, USA.

## Abstract

Microglia continuously monitor neuronal health through somatic purinergic junctions, where microglial processes establish dynamic contacts with neuronal cell bodies. The P2Y12 receptor is a key component of these junctions, essential for intercellular communication between ramified microglia and neurons under homeostatic conditions. While P2Y12 has long been considered a marker of homeostatic microglia, its potential role in reactive microglia during neurodegenerative disease remains largely unexplored. In this study, we demonstrate that P2Y12 deletion significantly reduces microglia-neuron process-to-body contacts in adult mice, consistent with previous findings. However, unexpectedly, P2Y12 loss markedly increases microglia-neuron body-to-body contacts, revealing an alternative mode of microglia-neuron communication independent of P2Y12. In prion-infected mice, P2Y12 expression persists in reactive, amoeboid microglia during advanced disease stages, including those engaging in extensive neuronal envelopment. Notably, P2Y12 loss increases the prevalence of envelopment events and accelerates disease progression. These findings redefine the role of P2Y12 in neurodegeneration, suggesting that its progressive decline lowers the threshold for microglia-neuron body-to-body interactions, ultimately influencing disease trajectory.

## Introduction

Microglia are the main cell type in the central nervous system (CNS) responsible for monitoring the brain environment and clearing injured or apoptotic cells. In their homeostatic state, microglia maintain neuronal surveillance by forming transient purinergic junctions between their processes and neuronal cell bodies ^1,2^. P2Y12 receptor, a member of the P2Y family of transmembrane purinoceptors that detects ATP and ADP, clusters on microglial processes at somatic junctions and plays a pivotal role in microglia-neuron communication ^1,2^. In addition to sensing neuronal health, P2Y12 mediates the elimination of compromised neurons or their components ^3-5^. By detecting ATP and ADP released by injured neurons, P2Y12 signaling facilitates the extension of microglial processes, their migration toward injury sites, and the engulfment of neuronal cell bodies and myelinated axons ^3-6^. Notably, P2Y12 signaling has been identified as essential for recruiting microglia to sites of CNS infection by neurotropic herpesviruses and for the phagocytosis of infected neurons ^5^.

In the brain, P2Y12 is exclusively expressed by resident microglia and is widely regarded as a marker of homeostatic microglia ^7-9^. While P2Y12 expression is robust under homeostatic conditions, it is significantly downregulated during sustained microglial activation in chronic neurological conditions, including neurodegenerative diseases ^9-14^. Notably, P2Y12 deficiency is considered as one of the features of damage-associated microglia in Alzheimer’s disease ^9^. Similarly, its expression is also downregulated in prion diseases ^15^. Not surprisingly, despite its critical role, P2Y12 has received relatively limited attention in the context of neurodegenerative diseases, leaving a significant gap in our understanding of its function in shaping the reactive microglial phenotype. How neuronal health is monitored under chronic neuroinflammation, when microglia adopt a reactive phenotype marked by reduced ramification and downregulation of P2Y12, is currently not known.

Prion diseases, also known as Transmissible Spongiform Encephalopathies, represent a class of fatal, transmissible neurodegenerative disorders affecting both humans and animals ^16^. The key pathological event underlying prion diseases involves the propagation and spread of the misfolded, disease-associated isoform of the prion protein (PrP^Sc^) throughout the CNS. This process involves the recruitment and structural conversion of the host’s normal cellular prion protein (PrP^C^) into the aberrant, β-sheet-rich isoform, PrP^Sc 17^. Neuronal degeneration in prion diseases is primarily attributed to the intrinsic neurotoxicity of PrP^Sc 18-24^. Unlike animal models of other neurodegenerative disorders, prion-infected animals develop genuine neurodegenerative disease, faithfully recapitulating key features of the human condition ^25^.

Microglia, as the primary cells responsible for monitoring neuronal health, are active participants in responding to neuronal dysfunction associated with prion infection. In fact, the progression of prion disease can be substantially modulated - either delayed or accelerated - through the manipulation of microglial phenotypes or their signaling pathways via genetic or pharmacological interventions ^26-32^. Our recent studies reveal that, starting in the late preclinical stages of prion disease, reactive microglia engage in a process of enveloping neuronal cell bodies, forming extensive body-to-body cell contacts ^33^. While this phenomenon superficially resembles engulfment, most neurons subjected to this process are only partially encircled by microglia and remain structurally intact. These neurons lack apoptotic markers but exhibit signs of functional decline. Partial neuronal envelopment was consistently observed across multiple prion-affected brain regions, in mice infected with diverse mouse-adapted prion strains, and in various subtypes of sporadic Creutzfeldt-Jakob disease in humans ^33^. However, the functional significance and pathological consequences of this process remain unclear ^34^.

Inspired by studies highlighting the critical role of P2Y12 in the engulfment and clearance of injured neurons, including those infected with neurotropic viruses ^5^, we investigated whether neuronal envelopment in prion disease relies on a P2Y12-dependent mechanism. We found that P2Y12 is expressed in reactive, amoeboid microglia, including those that envelop neurons in advanced stages of prion disease. In non-infected adult mice, P2Y12 deletion shifted microglia-neuron interactions from predominantly process-to-body contacts toward body-to-body contacts. In prion-infected mice, P2Y12 loss intensified neuronal envelopment by reactive microglia and accelerated disease progression. We speculate that body-to-body contacts represent an alternative, P2Y12-independent form of microglia-neuron interaction employed by reactive microglia to monitor neuronal health. Collectively, these findings redefine the role of P2Y12 in neurodegeneration, suggesting that its progressive decline may lower the threshold for body-to-body microglia-neuron interactions with chronic neurodegeneration, ultimately influencing disease outcomes.

## Results

### A subpopulation of reactive microglia retains P2Y12 expression in advanced stages of prion disease

P2Y12 is widely recognized as a marker of homeostatic microglia, with its expression being significantly downregulated in reactive microglia under neurodegenerative conditions ^7-9^. In prion-infected wild type (WT) mice, microglial activation follows PrP^Sc^ deposition in a spatially and temporally coordinated manner^19^. Unexpectedly, P2Y12 expression persists in late-stage disease, including the terminal stage (Fig. 1a, b). P2Y12 expression is observed in all prion-affected brain regions, including the thalamus, the area most severely affected (Fig. 1a-c; Fig. S1).

**Figure 1.**
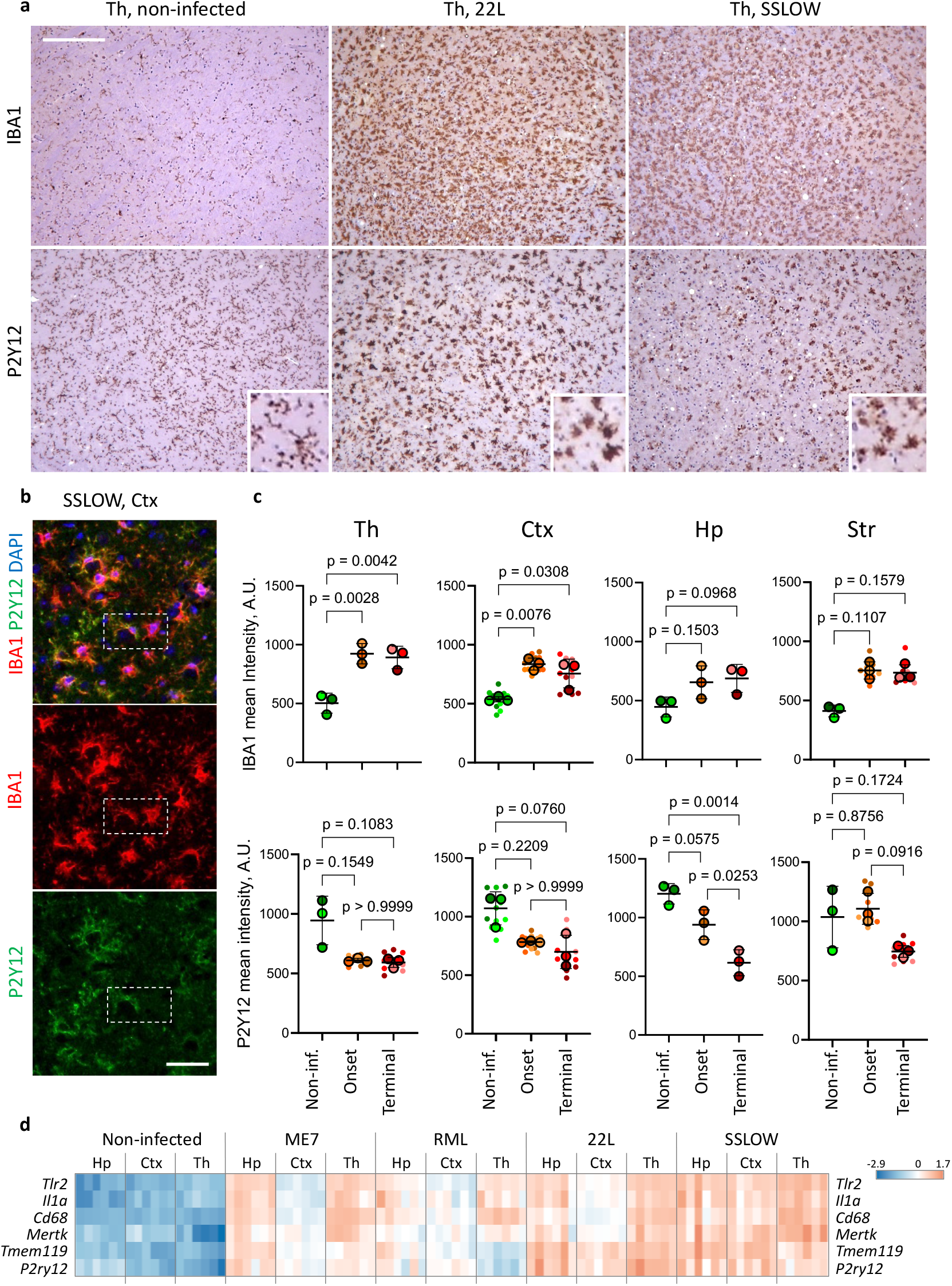
P2Y12 is expressed by reactive microglia in advanced stages of prion disease. **a**. Thalamus of non-infected WT mice and WT mice infected with 22L or SSLOW prion strains via i.c. stained for IBA1 (upper panel) or P2Y12 (lower panel) at the terminal stage of the disease. Magnified images in insets show P2Y12+ ramified or reactive microglia. Scale bar 100 µm. **b**. Cortex of WT mice infected with SSLOW via i.p. examined at terminal stage using co-immunostaining with anti--IBA1 (red) and anti-P2Y12 (green) antibodies. Scale bar 20 µm. Dashed rectangle highlights two cells with different P2Y12 intensity. **c**. Quantification of IBA1 and P2Y12 intensities in the thalamus (Th), cortex (Ctx), hippocampus (Hp), and striatum (Str) of WT mice infected with SSLOW via i.p. at the onset and terminal stage of the disease and non-infected WT control mice. Colors represent different brains. Dots represent individual fields of view. Average values for each brain are shown as circles. Means ± SD deviations are marked by black lines. N=3 brains per group, 1-6 images per brain region for each animal, Ordinary way ANOVA with Tukey’s multiple comparison for IBA1 in Th and Ctx, and P2Y12 in Hp and Str. Kruskal-Wallis multiple comparison for IBA1 in Hp and Str, and P2Y12 in Th and Ctx. **d**. Bulk expression of *P2ry12* at the terminal stage of the disease in thalamus (Th), cortex (Ctx), hippocampus (Hp), and striatum (Str) in WT mice infected with ME7, RML, 22L or SSLOW strains via i.p. route. Heatmap presents log10-transformed normalized counts obtained with NanoString.

Quantitative analysis of fluorescence microscopy images revealed region-specific differences in the dynamics of IBA1 and P2Y12 expression during disease progression (Fig. 1c). In all prion-affected regions, increased IBA1 expression was accompanied by a decline in P2Y12 levels in individual microglial cells (Fig. S1). Notably, in the thalamus, which is the most advanced with respect to the disease progression, IBA1 expression peaked while P2Y12 expression reached its lowest level by clinical onset. In contrast, these changes occurred more gradually in the cortex, hippocampus, and striatum, which exhibited less severe pathology (Fig. 1c).

Co-immunostaining of brains at the advanced disease stages for IBA1 and P2Y12 (Fig. 1b) highlighted heterogeneity in the microglial reactive states. Interestingly, both IBA1^+^/P2Y12^-^ and IBA1^+^/P2Y12^+^ myeloid cells exhibited the amoeboid morphology characteristic of reactive microglia, which starkly contrasted with the ramified morphology of homeostatic P2Y12^+^ microglia in uninfected animals (Fig. 1a, b). This P2Y12^+^ amoeboid microglial phenotype was consistently observed across different prion strains (Fig. 1a, and Fig. S1).

Despite the overall downregulation of P2Y12 in individual cells, bulk tissue gene expression analysis revealed higher levels of *P2ry12* expression at the terminal stage of the disease compared to control, non-inoculated animals (Fig. 1d). As P2Y12 expression is known to be strictly microglia-specific, the increased levels of *P2ry12* transcripts are attributed to microglial proliferation ^30,33,35^. The elevated *P2ry12* expression in bulk tissues was observed in all prion-affected brain regions and across all four prion strains analyzed: Me7, RML, 22L, and SSLOW (Fig. 1d).

In summary, P2Y12^+^ reactive microglia persist throughout the clinical stages of prion disease and remain detectable even at the terminal stage. In advanced disease stages, P2Y12 is expressed by reactive microglia with an amoeboid morphology.

### P2Y12^+^ microglia envelop neurons

Previously, we demonstrated that in prion disease, reactive microglia envelop neuronal cell bodies, forming extensive body-to-body contacts in a process superficially resembling neuronal engulfment ^33^. Given that P2Y12 signaling is implicated in the engulfment of injured neurons ^5^, we next investigated whether neuronal envelopment in prion-infected mice is carried out by P2Y12^+^ microglia. P2Y12^+^ microglia were observed in proximity to neuronal nuclei and displayed a cup-shaped, semicircular morphology, characteristics of reactive microglia engaged in neuronal envelopment (Fig. 2a). Confocal imaging confirmed direct body-to-body contacts between neuronal soma and reactive P2Y12^+^ microglia in SSLOW-infected mice (Fig. 2b, d). Consistent with previous studies, microglia involved in neuronal envelopment contained PrP^Sc^ deposits, suggesting that the P2Y12^+^ population participates in the phagocytic uptake of PrP^Sc^ (Fig. 2b-e). Next, we aimed at testing whether deletion of P2Y12 disrupts neuronal envelopment and influences disease progression.

**Figure 2.**
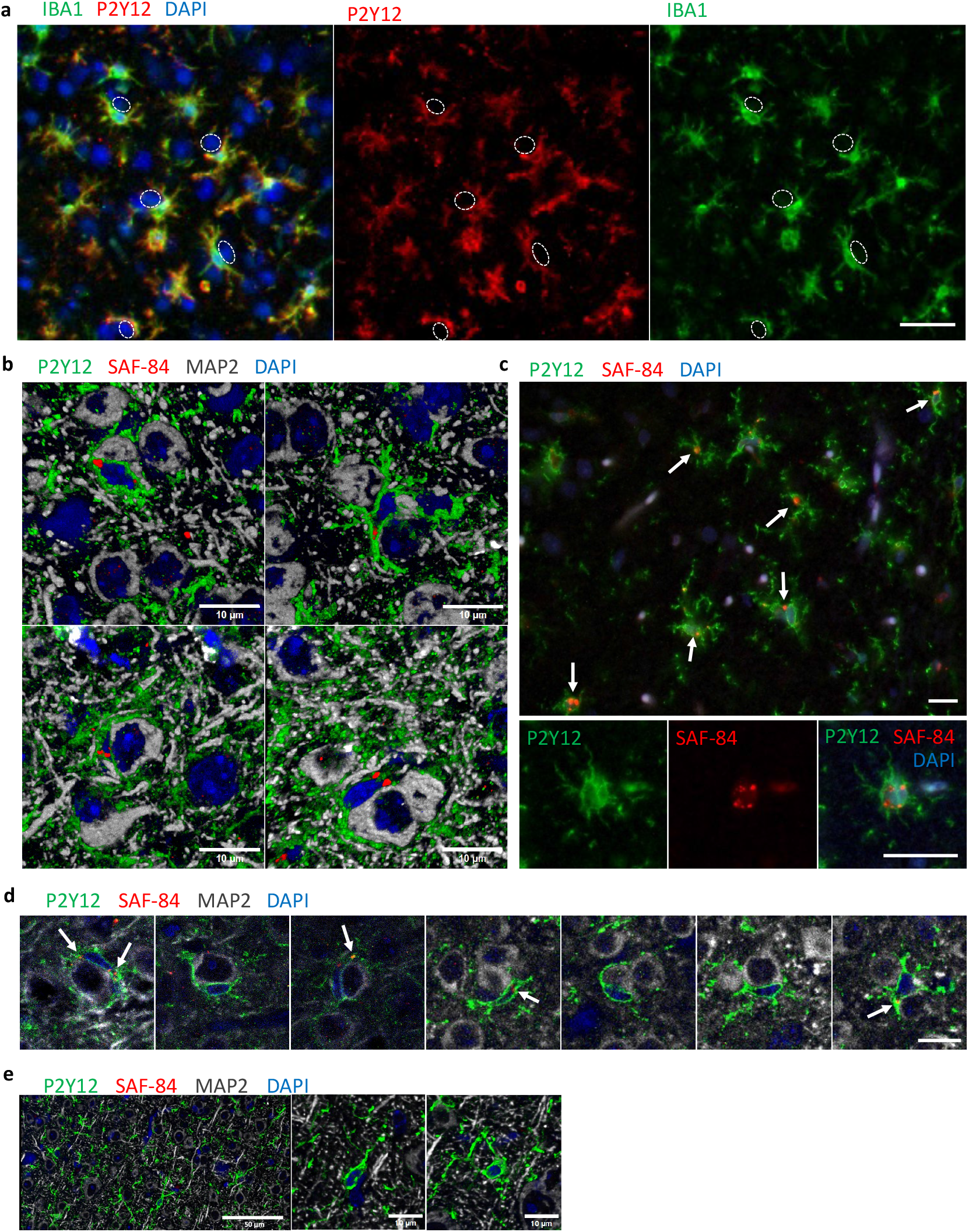
Neurons are enveloped by microglia positive for P2Y12. **a**. Co-immunofluorescence of IBA1 and P2Y12 in the cortex of SSLOW-infected mice showing cup-shaped microglia typical for envelopment. Dashed circles mark positions of neurons. **b**. Confocal 3D reconstruction images showing PrP^Sc^ (SAF-84, red) in P2Y12-positive microglia (green) enveloping neurons (MAP2, gray) in cortex of SSLOW-infected mice. **c**. Epifluorescence images of SSLOW-infected brains showing P2Y12-positive microglia (green) containing PrP^Sc^ (SAF-84, red). Arrows point to the examples of PrP^Sc^-containing P2Y12-positive microglia. **d**. A gallery of confocal images showing examples of PrP^Sc^ (SAF-84, red) in P2Y12-positive microglia (green) enveloping neurons (MAP2, gray). Arrows point to PrP^Sc^ puncta. **e**. Confocal 3D reconstruction images from cortex of non-infected brains co-immunostained for PrP^Sc^ (SAF-84, red), microglia (P2Y12, green), and neurons (MAP2, gray). Scale bars 20µm in **a**, 10 µm in **b, c** and **d**, 50 µm or 10 µm in **e**, as marked.

### P2Y12 deletion alters the morphology of homeostatic microglia in adult mice

Before assessing the effect of P2Y12 knockout (KO) on prion disease progression, we first examined whether its deletion influenced the homeostatic state of microglia. Morphological differences were observed between P2Y12 KO and wild-type (WT) microglia. Full-cortex scans analyzing 17 morphological parameters allowed division of IBA1^+^ cells into two sub-clusters, different with respect of the degree of ramification. Compared to WT mice, P2Y12 KO mice exhibited a higher proportion of less-ramified microglia (64.7 ± 2.9% in P2Y12 KO vs. 51.9 ± 1.4% in WT) and a lower proportion of more-ramified microglia (35.3 ± 2.9% in P2Y12 KO vs. 48.1 ± 1.4% in WT) (Fig. S2a,b).

Beyond morphological differences, P2Y12 KO microglia exhibited an upward trend in TMEM119 expression compared to WT microglia (Fig. S2c). Although TMEM119 is often regarded as a marker of homeostatic microglia, its transcripts are upregulated in the brains of Alzheimer’s disease patients ^36,37^.

Analysis of the IBA1^+^ cells revealed that, in P2Y12 KO mice, homeostatic microglia displayed higher circularity and solidity relative to WT microglia (Fig. 3a). Additionally, P2Y12 KO microglia showed a trend toward shorter branch lengths (Fig. 3a). These changes suggest a shift toward a more amoeboid morphology. Recent studies have highlighted sex-specific cellular perturbations in P2Y12 KO mice ^38^. Consistent with previous findings, we observed more pronounced morphological differences, particularly in circularity and solidity, between WT and P2Y12 KO microglia in females than in males (Fig. 3b). However, no statistically significant differences were detected between males and females within the same genotype, whether WT or P2Y12 KO (Fig. 3b).

**Figure 3.**
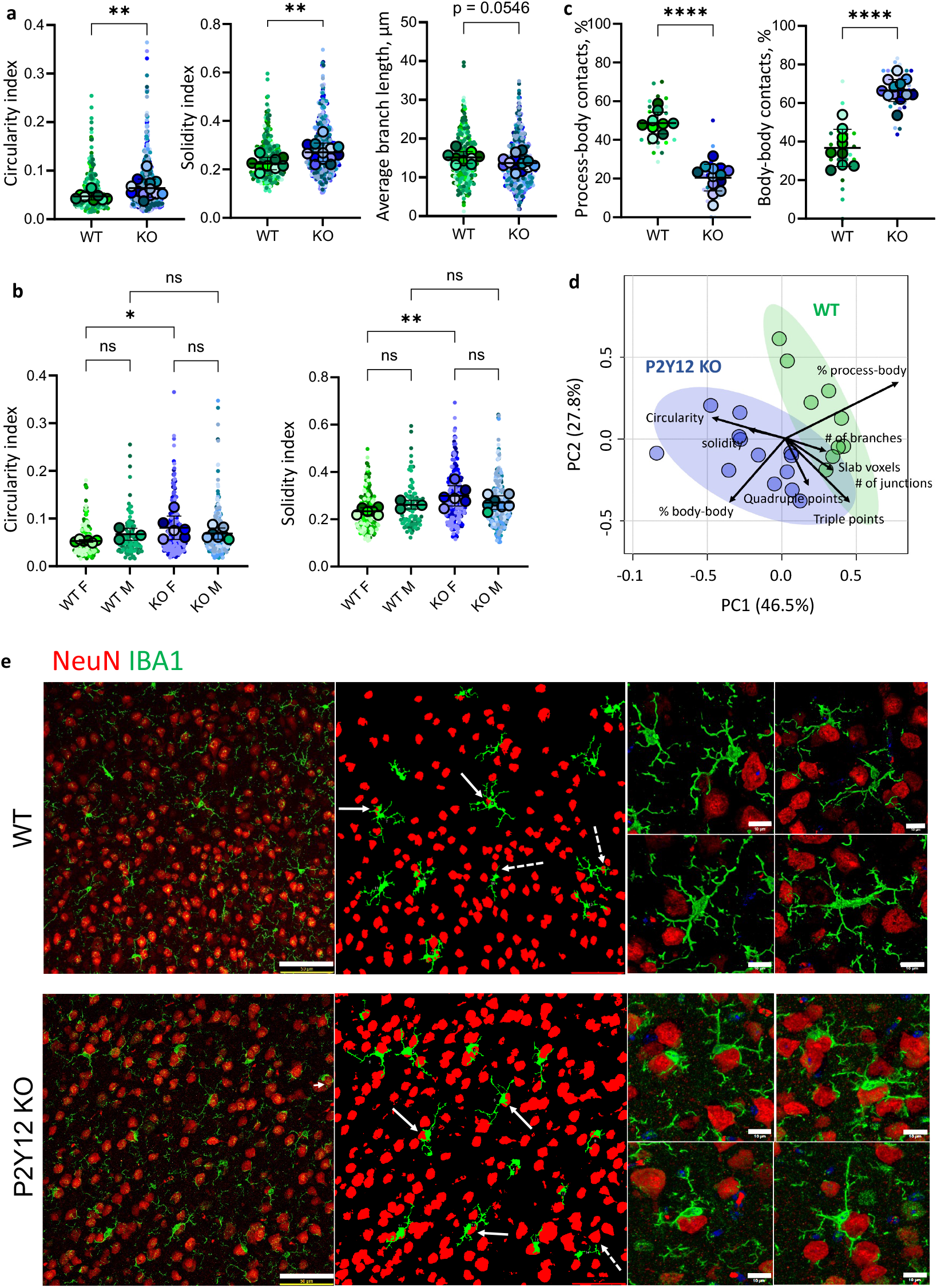
Analysis of microglia morphology in non-infected P2Y12 KO and WT mice. **a**. Comparison of P2Y12 KO and WT microglia by selected morphological parameters: circularity, solidity, and average branch length. **b**. Circularity and solidity data from **a** plotted for males and females separately. **c**. Quantification of process-body and body-body contacts between microglia and neurons in cortices of WT and P2Y12 KO mice. **d**. Principal component analysis of morphological parameters. In **a-d**, colors represent different brains; average values for each brain are shown as circles; dots represent individual cells in **a** and **b**, or fields of view in **c**. Means ± SD are marked by black lines. In **a, b**, and **c**, *p<0.05, **p<0.01, ****p<0.0001, ns – non-significant. For circularity, medians were compared by unpaired t-test with Welch’s correction. For solidity, medians were compared by unpaired Student’s t-test. For average branch length, process-body and body-to-body contacts, means were compared by unpaired Student’s t-test, n=9-13 brains for each group, 2-4 fields of view per brain. In **b**, circularity and solidity compared by ordinary ANOVA with Tukey’s multiple comparison, n=3-7 brains for each group. **e**. Imaging of process-to-body and body-to-body contacts between microglia (IBA1, green) and neurons (NeuN, red). 3D maximum intensity projections images are shown on the left, corresponding merged binary images – in the middle, and magnified 3D reconstruction images of microglia in contact with neurons are shown on the right. Dashed and solid arrows indicate process-to-body and body-to-body contacts, respectively. Scale bars 50 μm (left) and 10 μm (right).

In conclusion, our findings indicate that P2Y12 deletion alters the morphology of homeostatic microglia, specifically reducing their degree of ramification. These results align well with prior research^38,39^.

### Deletion of P2Y12 promotes microglial-neuronal body-to-body contacts

To investigate whether P2Y12 influences microglia-neuron interactions, we analyzed two types of contacts using confocal microscopy: microglial process-to-neuronal body (process-to-body) and microglial body-to-neuronal body (body-to-body) contacts (Fig. 3e). In P2Y12 KO mice, the percentage of IBA1+ microglia exhibiting process-to-body contacts with neurons was significantly lower than in WT mice (Fig. 3c, e). Conversely, the proportion of IBA1+ microglia forming body-to-body contacts was substantially higher in P2Y12 KO compared to WT mice (Fig. 3c, e). This suggests that the increase in body-to-body contacts in P2Y12 KO mice occurs at the expense of process-to-body interactions. Microglia engaged in body-to-body contacts retained a predominantly ramified morphology, indicative of a homeostatic phenotype (Fig. 3e).

Principal component analysis (PCA) of microglial morphology and microglia-neuron contacts revealed a clear separation between P2Y12 KO and WT populations, highlighting shifts in the homeostatic state of microglia associated with P2Y12 loss (Fig. 3d). The primary drivers of this separation, which distinguished the P2Y12 KO and WT genotypes, were the differences in contact type along with circularity. In conclusion, P2Y12 deletion alters microglia-neuron interactions, shifting the contact type from process-to-body toward body-to-body.

### P2Y12 deletion accelerates prion disease progression after clinical onset

To examine the impact of P2Y12 deletion on disease progression, P2Y12 KO and WT mice were intracerebrally (i.c.) inoculated with a mouse-adapted prion strain SSLOW ^40,41^. In SSLOW-infected WT mice, clinical onset typically occurs around 95 days post-inoculation (dpi), while terminal disease, defined as a 20% loss in body weight after a progressive worsening of neurological symptoms, is reached at 123 ± 4 dpi (mean ± SD) ^41^. Clinical disease, assessed by a combined score assessing posture, mobility, clasping, and gait abnormalities, developed simultaneously in both SSLOW-infected P2Y12 KO and WT cohorts (Fig. 4a). However, following clinical onset, disease progression was significantly accelerated in P2Y12 KO mice (Fig. 4b). Consequently, mean survival was markedly shorter in P2Y12 KO mice compared to WT controls (mean ± SD: 111.3 ± 4.4 dpi vs. 121.8 ± 5.1 dpi, respectively) (Fig. 4c). This survival difference was consistent across both male and female cohorts (Fig. S3a).

**Figure 4.**
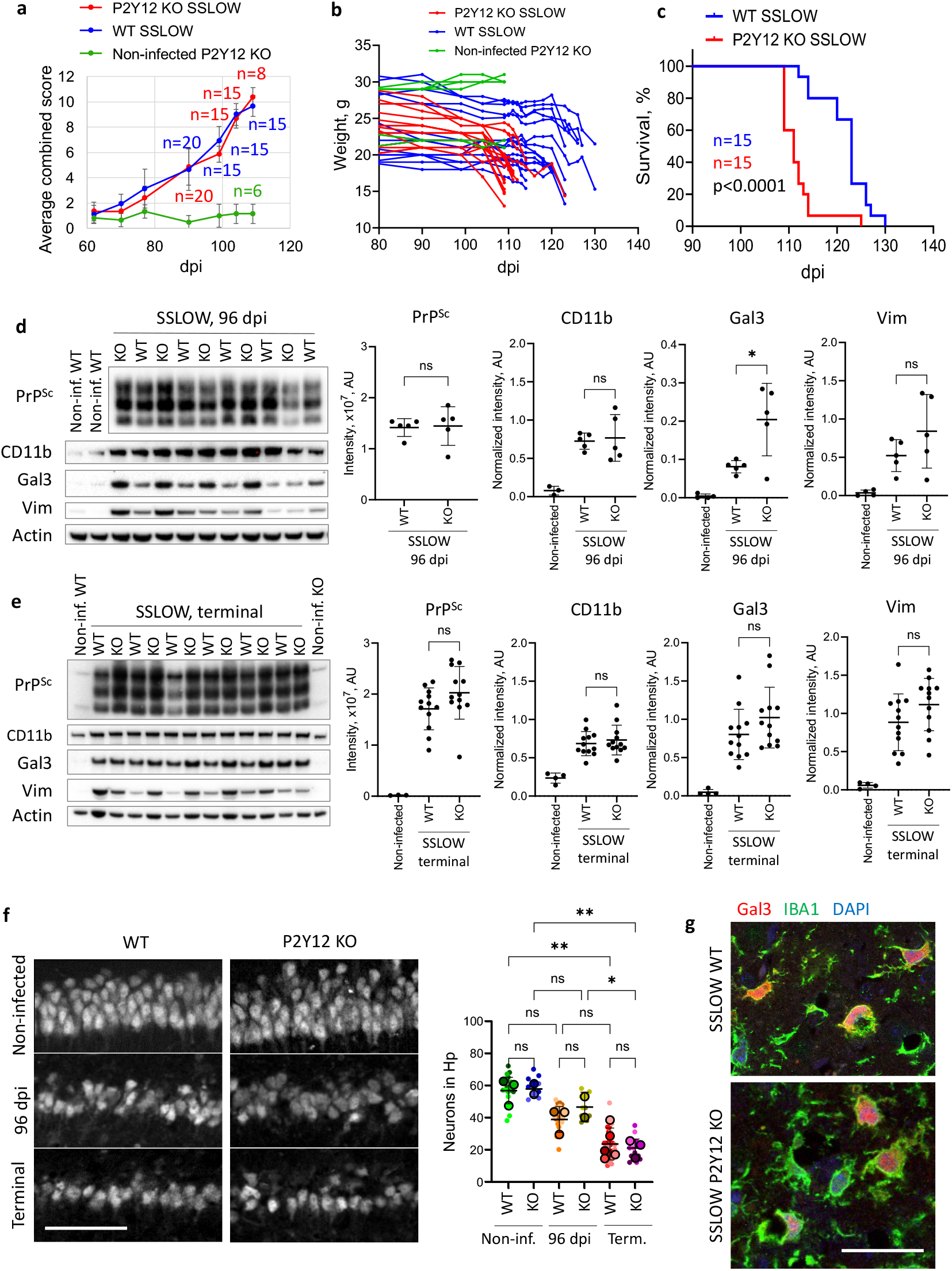
P2Y12 deletion accelerates prion disease progression. P2Y12 KO and WT mice were inoculated with SSLOW via i.c. route and monitored for disease progression. **a** and **b**. Progression of clinical signs, as monitored by scoring of the clasp, posture, mobility, and gait, and presented as averaged combined scores ± SD (**a**), and weight changes in individual SSLOW-infected P2Y12 KO and WT mice (**b**). Non-infected age-matched P2Y12 KO mice are provided as reference. **c**. Survival curves for SSLOW-infected P2Y12 KO and WT mice. Comparison by Mantel-Cox test, n=15 for each group. **d** and **e**. Representative Western blot images and quantification of selected markers in brains of SSLOW-infected P2Y12 KO and WT mice at 96 dpi (**d**) and the terminal stage (**e**). The data presented as Means ± SD. *p<0.05, ns - non-significant, by unpaired Student’s t-test, n=5 per group at the 96 dpi, and n=12 per group at the terminal stage. Data for non-infected brains are shown as a reference. **f**. Loss of pyramidal neurons in CA1 area of hippocampus. Left: representative immunofluorescence images of neurons in P2Y12 KO and WT stained for NeuN in non-infected brains, at 96 dpi, and at the terminal stages of the disease. Right: analysis of neuronal density by ordinary one-way ANOVA with Tukey’s multiple comparison test, n=2-5 animals in each group, 3-12 measurements per brain. Colors represent different brains; dots represent individual neuronal counts taken from 100×200 pixels rectangles within CA1 pyramidal layer. Average values for each brain are shown as circles. Means ± SD are marked by black lines. *p<0.05, **p<0.01, ns non-significant. **g**. Confocal images of SSLOW-infected WT (upper panel) and P2Y12 KO (lower panel) cortices co-stained for Gal3 (red) and IBA1 (green). Scale bars 50 µm in **f**, 20 µm in **g**.

To investigate potential pathological mechanisms underlying the accelerated disease progression in P2Y12 KO mice, we assessed PrP^Sc^ accumulation, neuronal loss in the hippocampal pyramidal layer, and astrocyte reactivity. No significant differences were observed between SSLOW-infected P2Y12 KO and WT mice in PrP^Sc^ levels at either clinical onset or terminal disease stages (Fig. 4d, e). Similarly, total PrP (PrP^C^ + PrP^Sc^) levels remained comparable between genotypes at both disease stages (Fig. S3b, c). Both genotypes exhibited comparable neuronal loss in the hippocampal pyramidal layer at either clinical onset or terminal stage (Fig. 4f). Lastly, astrocyte reactivity, assessed via Vim expression, showed no significant differences between genotypes at either stage of the disease (Fig. 4d, e).

### Differential dynamics of Gal3 upregulation in SSLOW-infected P2Y12 KO and WT mice

To evaluate the impact of P2Y12 deletion on microglial phenotype in SSLOW-infected mice, we analyzed several markers, including CD11b, IBA1, TMEM119, and Gal3 along with the gene expression of purinergic receptors (*P2ry13, P2ry6, P2rx4*, and *P2rx7*). Immunofluorescence microscopy confirmed the loss of P2Y12 in P2Y12 KO mice (Fig. 5a, b). Western blot analysis revealed no significant differences in CD11b expression between SSLOW-infected P2Y12 KO and WT mice at either the clinical onset or terminal stages of the disease (Fig. 4d, e). Immunostaining revealed increased IBA1 expression in both SSLOW-infected P2Y12 KO and WT mice compared to their respective non-infected controls, though signal intensity remained comparable between the infected groups (Fig. 5a, b). Similarly, TMEM119 immunofluorescence intensity did not differ between SSLOW-infected P2Y12-KO and WT mice (Fig. 5c).

**Figure 5.**
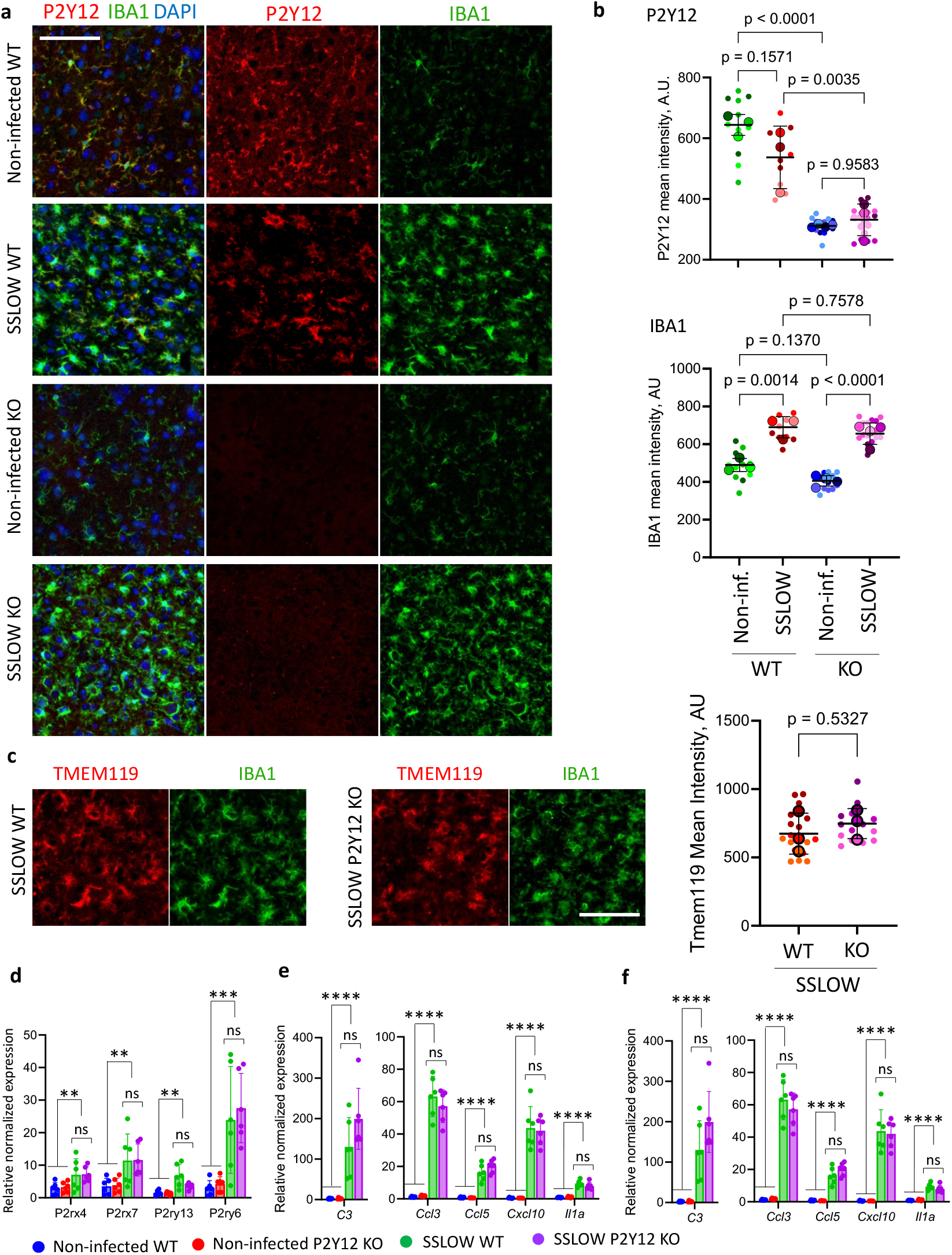
Analysis of microglial markers in SSLOW-infected P2Y12 KO and WT mice. **a**. Immunofluorescence staining for P2Y12 (red) and IBA1 (green) showing lack of P2Y12 signal in non-infected and SSLOW-infected P2Y12 KO brains. Cortex is shown. **b**. Quantification of P2Y12 and IBA1 signal intensities in cortices of non-infected and SSLOW-infected WT and P2Y12 KO mice. Comparison by ordinary one-way ANOVA with Tukey’s multiple comparison test, n=3-4 animals in each group, 3-6 fields of view per brain. **c**. TMEM119 (red) and IBA1 (green) co-immunostaining (left) and TMEM119 quantification (right) in the cortices of SSLOW-infected WT and P2Y12 KO mice. Comparison by unpaired Student’s t-test, n=3 brains for each group, 3-7 fields of view per brain. In **b** and **c**, colors represent different brains, dots represent fields of view, average values for each brain are shown as circles. Means ± SD are marked by black lines. **d**,**e**,**f**. RT-qPCR analysis of selected purinergic receptors enriched in microglia (**d**), neuroinflammatory markers (**e**), and markers of phagocytic activity (**f**). Comparison of non-infected WT, non-infected KO, SSLOW WT and SSLOW KO was performed using Brown-Forsythe ANOVA followed by Dunnett’s T3 multiple comparisons test, n=6 brains per group; ns – not significant; non-infected WT vs KO are not shown (not shown). Comparison of combined non-infected WT+KO vs combined SSLOW WT+KO was performed using unpaired two-tailed t-test with Welch’s correction for *C3, Ccl3, Ccl5, Il1a, Tmem119*, Cd68, *Trem2, Tlr2, P2rx4, P2rx7, P2ry13, P2ry6*, and non-parametric two-tailed Mann-Whitney test for *Cxcl10 and Mertk*; n=12 brains per combined group; **p<0.01, ***p<0.001, ****p<0.0001. Scale bars 50 µm.

Analysis of gene expression of the purinergic receptors (*P2rx4, P2rx7, P2ry13* and *P2ry6*) showed upregulation in SSLOW-infected animals compared to non-infected controls, however, no differences between prion-infected P2Y12 KO and WT groups were found (Fig. 5d). Analysis of microglia-enriched genes associated with neuroinflammation (*C3, Il1a, Ccl3, Ccl5*, and *Cxcl10*) also revealed significant upregulation in SSLOW-infected groups, with no differences between P2Y12 KO and WT mice (Fig. 5e). Similarly, genes linked to phagocytic activity (*Trem2, Mertk, Cd68*, and *Tlr2*) were upregulated in infected groups, but no significant differences were observed between them (Fig. 5f).

Microglial morphology analysis revealed subtle differences between SSLOW-infected P2Y12 KO and WT mice, with P2Y12 KO microglia displaying a smaller cell area and perimeter than WT microglia (Fig. S3e). These morphological changes suggest a more advanced amoeboid phenotype in P2Y12 KO microglia. PCA of morphological parameters further confirmed these differences, showing a separation between P2Y12 KO and WT clusters (Fig. S3f).

Gal3 expression, quantified by Western blot, was significantly upregulated in both SSLOW-infected P2Y12 KO and WT mice compared to non-infected controls (Fig. 4d, e). Notably, by 96 dpi, Gal3 levels were higher in SSLOW-infected P2Y12 KO mice compared to WT counterparts (Fig. 4d). However, by the terminal stage, Gal3 expression in WT group had caught up to the P2Y12 KO level (Fig. 4e). Gal3 is known to play a critical role in lysosomal repair and is upregulated in response to lysosomal leakage ^42,43^. Consistent with this function, immunofluorescence microscopy confirmed the intracellular localization of Gal3 within IBA1^+^ myeloid cells in both SSLOW-infected P2Y12 KO and WT mice (Fig. 4g, and Fig. S3d). These findings suggest that microglia were the primary sites of lysosomal damages, which are likely attributed to the uptake and accumulation of PrP^Sc^. Notably, Gal3 level increased more rapidly in SSLOW-infected P2Y12 KO mice compared to WT mice. Overall, the phenotypic differences between reactive microglia in P2Y12 KO and WT mice appear to be subtle, with the primary distinction being the dynamics of Gal3 upregulation.

### Differential dynamics of PrP^Sc^ uptake in P2Y12 KO and WT mice

Given the differential dynamics of Gal3 upregulation, we next evaluated PrP^Sc^ uptake by microglia. While microglia do not replicate PrP^Sc^, they can acquire PrP^Sc^ positivity through phagocytic uptake ^33,44,45^. Previously, we demonstrated that microglia effectively phagocytose PrP^Sc^ during the preclinical stages. However, as the disease progresses into the clinical stages, microglial activity shifts toward establishing extensive neuron-microglia body-to-body contacts. ^33^.

Accumulation of PrP^Sc^ in IBA1^+^ cells was quantified using confocal microscopy imaging (Fig. 6a, b). At the onset of clinical disease, the number of PrP^Sc^-positive microglia was higher in P2Y12 KO mice than in WT mice (Fig. 6c). However, by the terminal stage, the number of PrP^Sc^-positive microglia in WT mice had caught up to that in P2Y12 KO mice (Fig. 6c). This pattern closely parallels the dynamics of Gal3 upregulation, reinforcing the link between PrP^Sc^ accumulation and lysosomal dysfunction.

**Figure 6.**
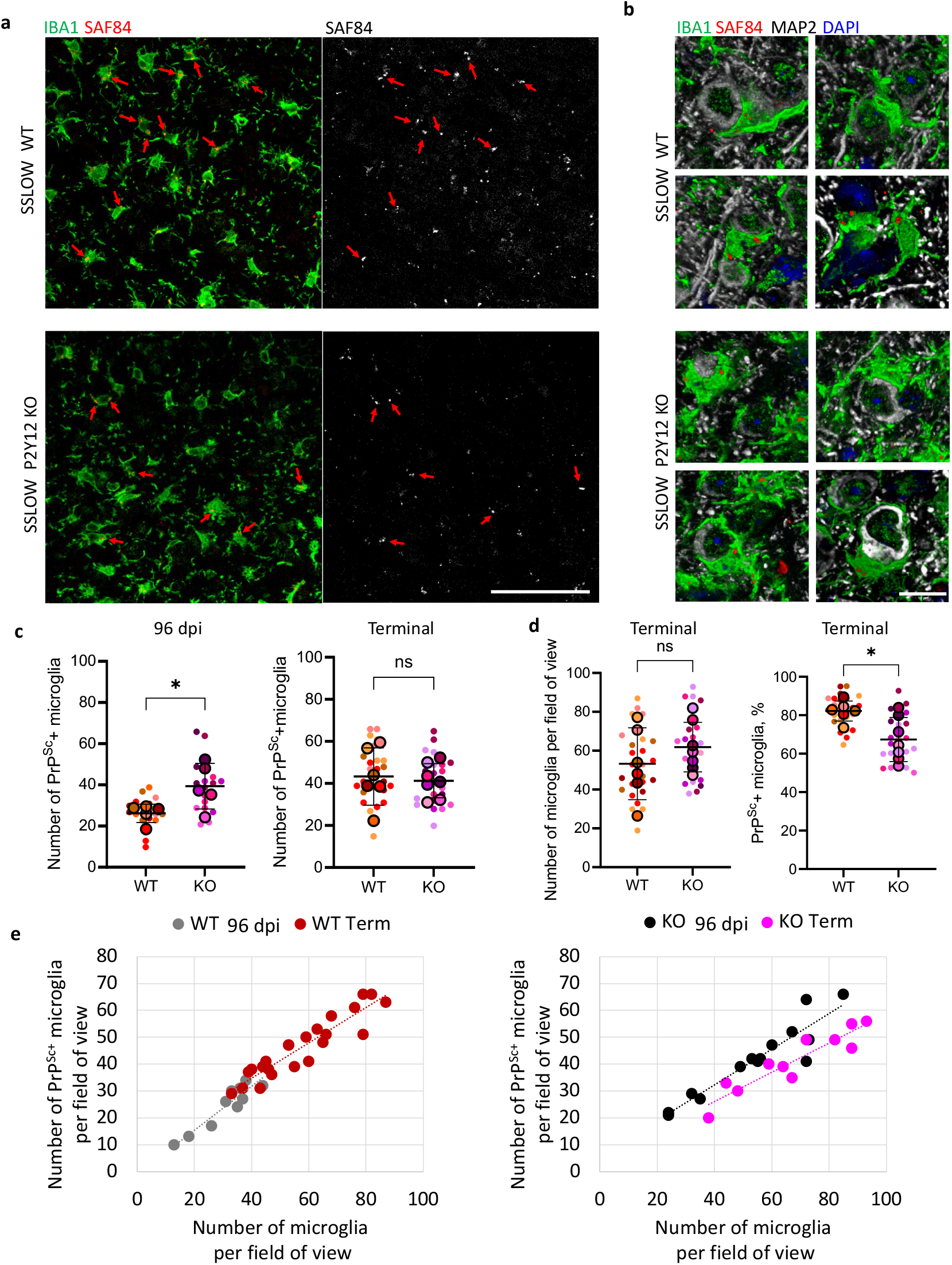
Differential dynamics of PrP^Sc^ uptake by IBA1^+^ cells in P2Y12 KO and WT mice. **a**. Confocal microscopy images of brains co-immunostained for PrP^Sc^ (SAF-84) and microglia (IBA1) and showing localization of PrP^Sc^ puncta to IBA1^+^ cells (left) in cortices of SSLOW-infected WT (top) and P2Y12 KO mice (bottom). SAF-84 channel images are shown in gray scale for clarity (right). Red arrows point at PrP^Sc^ puncta in IBA1^+^ cells. Scale bar 50 µm. **b**. 3D confocal microscopy images of cortices of SSLOW-infected WT (top) and P2Y12 KO (bottom) mice showing PrP^Sc^ puncta (SAF-84, red) localized to IBA1^+^ cells (green) enveloping neurons (MAP2, gray). Scale bar 10 µm. **c**. Quantification of PrP^Sc+^ IBA1^+^ cells in the cortices of SSLOW-infected WT and P2Y12 KO mice at 96 dpi and terminal stage of the disease. **d**. Quantification of IBA1^+^ cells per field of view and percent of PrP^Sc+^ IBA1^+^ cells at the terminal stage of the disease. In **c** and **d**, colors represent different brains; dots represent fields of view; average values for each brain are shown as circles. Means ± SD are marked by black lines. *p<0.05, ns – not significant by unpaired Student’s t-test, n=5-7 brains for each group, 2-8 fields of view per brain. **e**. Relationships between the number of IBA1^+^ cells and number of PrP^Sc+^ IBA1^+^ cells per field of view in WT (left) and P2Y12 KO (right) brains infected with SSLOW and analyzed at 96 dpi or terminal stage. Data points represent individual fields of view obtained from 5-7 cortices. Dashed lines represent linear trendline for each group.

The total density of IBA1^+^ cells, as quantified at the terminal stage, was comparable between WT and P2Y12 KO mice, with a modest uptrend in P2Y12 KO mice (Fig. 6d). Notably, when PrP^Sc^-positive IBA1^+^ cells were expressed as a percentage of total IBA1^+^ cells, P2Y12 KO mice exhibited a lower proportion of PrP^Sc^-positive IBA1^+^ cells than WT mice at the terminal stage (Fig. 6d). This suggests a more significant shift in myeloid cell activity away from PrP^Sc^ uptake in P2Y12 KO versus WT mice at the final stage of the disease. Indeed, plotting the number of PrP^Sc^-positive IBA1^+^ cells against the total number of IBA1^+^ cells per field of view reveals distinct dynamics in cell activity between the two groups. In SSLOW-infected WT mice, the number of PrP^Sc^-positive IBA1^+^ cells continues to rise at a rate proportional to total number of IBA1^+^ cells from clinical onset to the terminal stage, forming a single linear continuum (Fig. 6e). This trend suggests that WT IBA1^+^ cells continue accumulating PrP^Sc^ at a consistent rate until the terminal stage. In contrast, in P2Y12 KO, IBA1^+^ cells exhibited an altered uptake pattern. By 96 dpi, the number of PrP^Sc^-positive cells in P2Y12 KO mice was already high and comparable to that of terminal-stage WT mice (Fig. 6e). However, as the disease progressed, the increase in total P2Y12 KO IBA1^+^ cells was not accompanied by a corresponding rise in PrP^Sc^-positive microglia, indicating a loss of PrP^Sc^ uptake capacity (Fig. 6e).

In conclusion, the dynamics of Gal3 upregulation closely mirror the pattern of PrP^Sc^ uptake. P2Y12 KO mice reach their maximum PrP^Sc^ uptake capacity earlier than WT mice. While WT IBA1^+^ cells continue to uptake PrP^Sc^ after clinical onset, the rate of PrP^Sc^ uptake in P2Y12 KO slows considerably during clinical stage.

### P2Y12 deletion facilitates neuronal envelopment

Considering that P2Y12 deletion promotes microglia-neuron body-to-body contacts under normal conditions, we next evaluated whether P2Y12 depletion facilitates neuronal envelopment in SSLOW-infected mice. We quantified the percentage of neurons undergoing envelopment in combined cortical layers 2 to 6, which exhibited prion deposition (Fig. 7a, b). Consistent with previous studies ^2^, a modest downward trend in neuronal density within this area was observed in P2Y12 KO mice compared to WT mice, whether infected or non-infected (Fig. 7c). The percentage of neurons undergoing envelopment by microglia at clinical onset showed an upward trend in P2Y12 KO mice relative to WT mice (Fig. 7d). By the terminal stage, neuronal envelopment was significantly higher in P2Y12 KO mice than in WT mice (Fig. 7d). These findings suggest that P2Y12 deletion not only promotes body-to-body contacts between homeostatic microglia and neurons but also facilitates the envelopment of neurons by reactive microglia.

**Figure 7.**
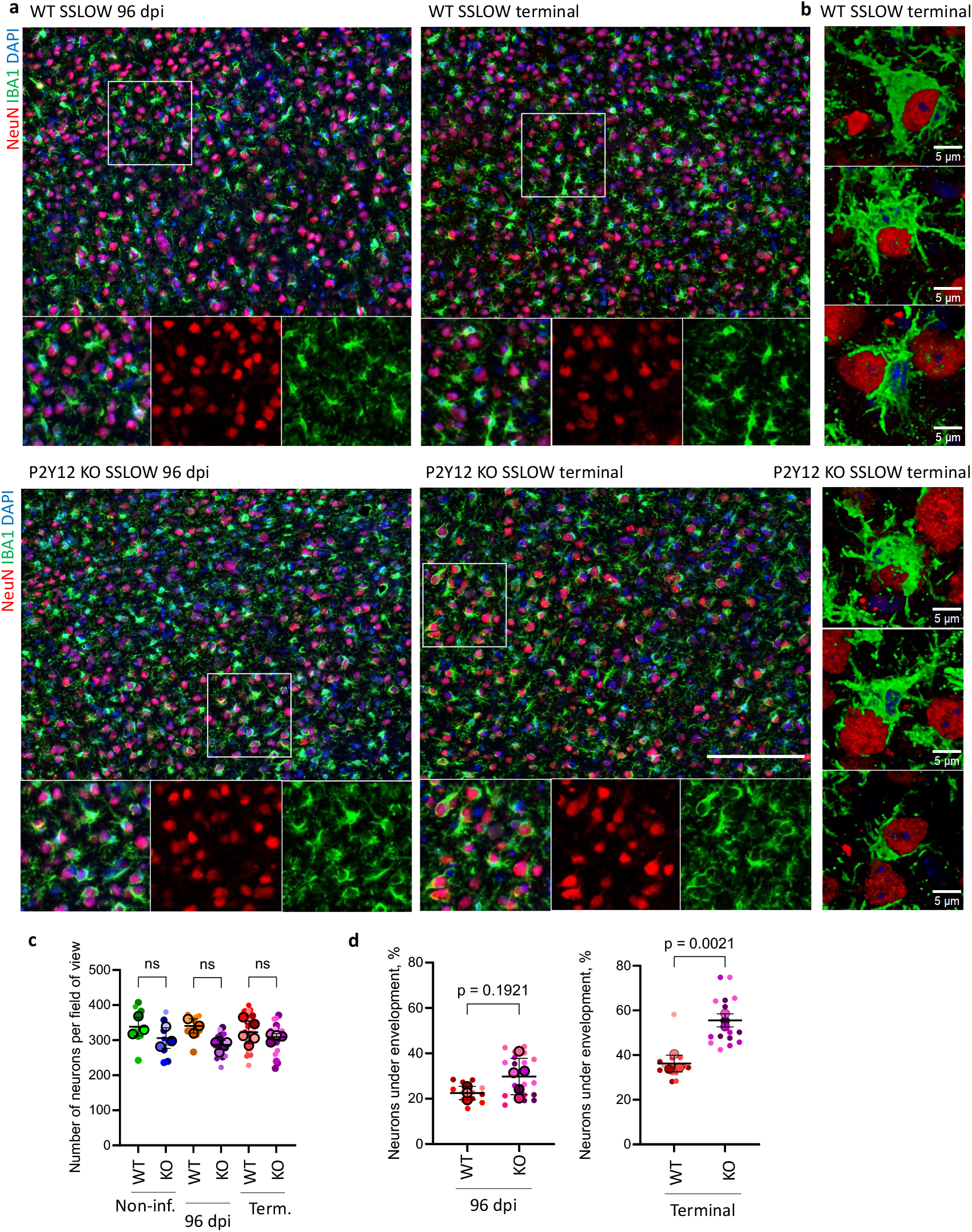
P2Y12 deletion facilitates neuronal envelopment in SSLOW-infected mice. **a**. Immunostaining of SSLOW-infected brains for microglia (IBA1, green) and neurons (NeuN, red) showing envelopment in WT (top) and P2Y12 KO cortices (bottom) at 96 dpi (left) and terminal stage of the disease (right). Insets present magnified images as merged and separated channels. Scale bar 100 µm. **b**. 3D confocal images showing examples of neuronal envelopment (NeuN, red) by microglia (IBA1, green) in WT (top) and P2Y12 KO mice (bottom). Scale bar 5 µm. **c**. Quantification of neurons per field of view in cortical layers 2 to 6 of non-infected and SSLOW-infected WT and P2Y12 KO mice at 96 dpi, and at the terminal stage. **d**. Quantification of neuronal envelopment in the cortices of SSLOW-infected WT and P2Y12 KO mice at 96 dpi (left) and terminal stage (right). In **c** and **d**, colors represent different brains; dots represent individual fields of view; average values for each brain are shown as circles. n=3-5 brains per group with 3-7 fields of view for each brain. Means ± SD are marked by black lines. ns - non-significant by ordinary one-way ANOVA with Tukey’s multiple comparison in **c**. Student’s t-test was used for statistical analysis in **d**.

### P2Y12 deletion suppresses basal motility of reactive microglia

Assuming that reactive microglia monitor neuronal health by forming extensive body-to-body contacts with one neuron at a time before moving to the next, a reduction in microglia motility would prolong these contacts at the expense of overall movement. P2Y12 signaling is critical for the migration of homeostatic microglia ^3^. However, whether the P2Y12 receptor is important for the basal motility of reactive microglia remains unknown.

To address this question, microglia were acutely isolated using CD11b-beads from SSLOW-infected P2Y12 KO and WT mice, along with non-infected age-matched control groups. Basal microglial motility was assessed in culture in the absence of external stimulation. As expected, in non-infected mice, both WT (Supplementary Video 1) and P2Y12 KO microglia (Supplementary Video 2) exhibited minimal motility, as indicated by low track speed and short distances covered (Fig. 8a-c). In contrast, reactive microglia from SSLOW-infected WT mice displayed a broad distribution in motility parameters, with the majority of cells exhibiting high speed and extensive movement (Fig. 8a-c, Supplementary Video 3). Reactive microglia from SSLOW-infected P2Y12 KO mice also demonstrated greater motility compared to non-infected controls (Supplementary Video 4). However, relative to their SSLOW-infected WT counterparts, a significant shift toward reduced speed and shorter travel distances was observed in P2Y12 KO reactive microglia (Fig. 8a-c).

**Figure 8.**
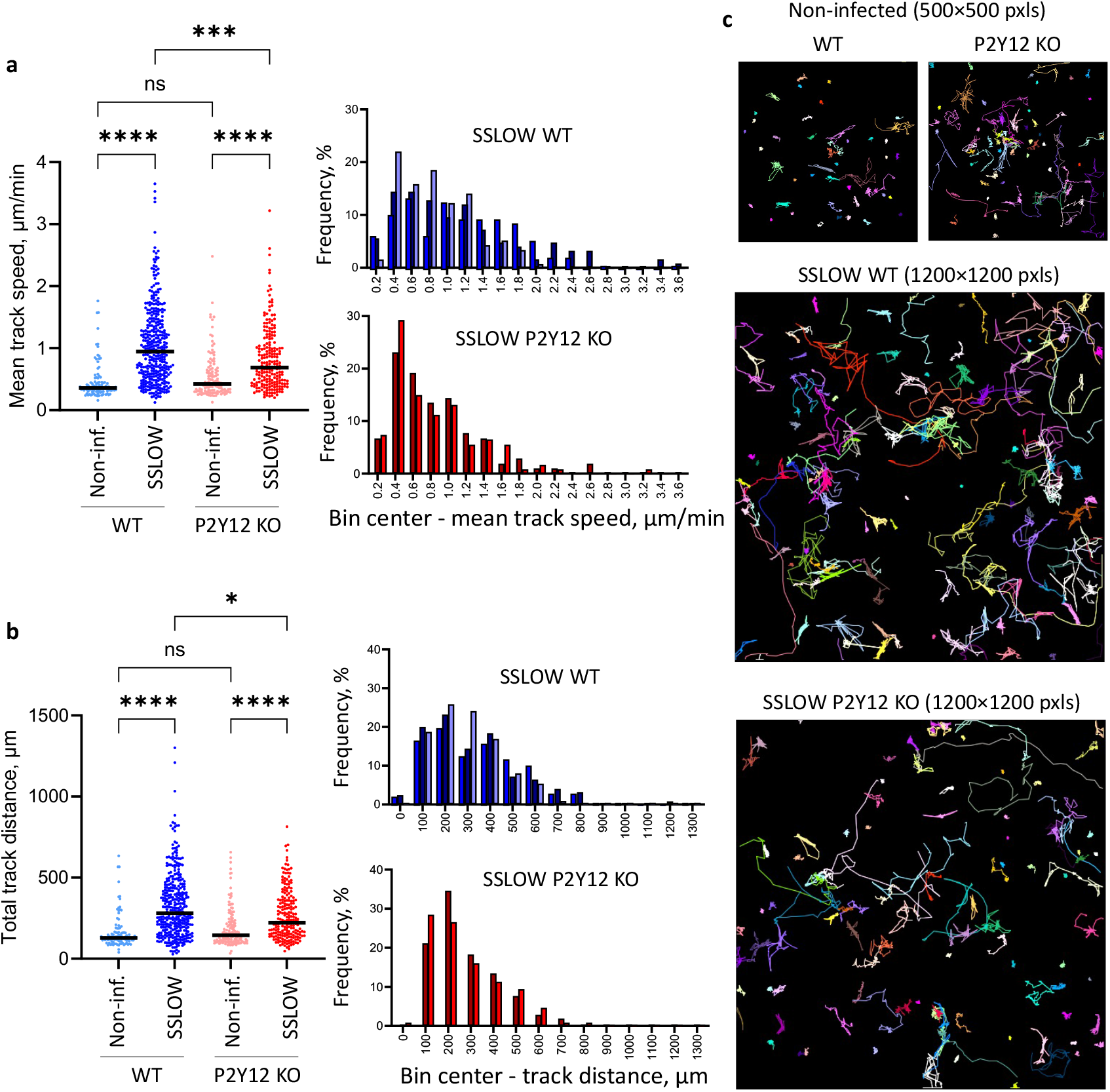
P2Y12 deletion suppresses basal motility of reactive microglia. CD11b^+^ cells were isolated at the clinical onset of the disease from WT and P2Y12 KO mice infected with SSLOW via i.p route. **a, b**. Mean track speed (**a**) and total track distance (**b**) of microglia isolated from non-infected and SSLOW-infected cortices of WT and P2Y12 KO mice and plotted as individual data points (left) or histograms (right). In **a** and **b**, *p<0.05, ***p<0.001, ****p<0.0001, ns – non-significant Kruskal-Wallis multiple comparison test, 106 – 361 cells per group. **c**. Examples of tracks recorded from non-infected and SSLOW-infected WT and P2Y12 KO primary microglia. Colored lines represent tracks of individual cells recorded within 6-hour period.

In conclusion, reactive microglia from prion-infected clinically ill animals exhibit high basal motility *in vitro*, even in the absence of external stimuli. However, P2Y12 deletion significantly impairs this basal motility. These findings support the hypothesis that the increased prevalence of neuronal envelopment in SSLOW-infected P2Y12 KO animals may result from reduced basal mobility of reactive microglia.

### The effects of P2Y12 deletion in mice infected with 22L prion strain are similar to those of SSLOW-infected mice

To determine whether the impact of P2Y12 deletion is strain-specific, we examined disease progression in mice infected with the 22L mouse-adapted prion strain. Clinical signs, assessed using the same composite score, developed in parallel in both 22L-infected P2Y12 KO and WT cohorts. Similar to SSLOW-infected mice, disease progression accelerated following clinical onset in 22L-infected P2Y12 KO mice compared to WT mice (Fig. S4a). Consequently, 22L-infected P2Y12 KO mice succumbed to prion disease faster than their WT counterparts. The amounts of PrP^Sc^ or total PrP (PrP^Sc^+PrP^C^) did not differ between P2Y12 KO and WT groups (Fig. S4b). However, the percentage of neurons under envelopment was significantly higher in 22L-infected P2Y12 KO compared to WT mice, repeating the pattern observed in SSLOW-infected cohorts (Fig. S4a).

## Discussion

Microglia continuously monitor neuronal activity and health via somatic purinergic junctions, formed by microglial processes in contact with neuronal bodies ^1^. As a key component of these junctions, the P2Y12 receptor is essential for intercellular communication between microglia and neurons ^1,2^. Under homeostatic conditions, process-to-body contacts serve as the primary mode of microglia-neuron interaction. Given that purinergic junctions rely on P2Y12, it is not surprising that P2Y12 deletion significantly reduced the frequency of process-to-body contacts in the present study, consistent with previous findings in P2Y12-deficient mice ^1^. Unexpectedly, however, the loss of P2Y12 significantly increased body-to-body contacts, suggesting an alternative mode of microglia-neuron communication. Unlike process-to-body interactions, body-to-body contacts are independent of P2Y12, but may involve alternative purinergic receptors.

P2Y12 is known to regulate microglial responses to acute injuries and shape their function during neurodevelopment. During CNS development, P2Y12 signaling is crucial for the engulfment and clearance of apoptotic cells ^46^. In response to neurotrophic viral infections, P2Y12 mediates microglial recruitment to infection sites and facilitates the phagocytosis of infected neurons ^5^. Similarly, in localized axonal injuries, microglia employ P2Y12-dependent protective wrapping to prevent axonal degeneration ^47^. Furthermore, in neuropathic pain models, microglia engulf both injured and uninjured myelinated axons in a P2Y12-dependent manner ^4^. These studies established that in acute injury and infection, P2Y12 signaling, together with a highly ramified morphology, enables homeostatic microglia to fulfill their surveillance functions. However, how microglia perform these critical tasks upon transitioning into a reactive, ameboid state remains unclear. In persistently activated microglia, both ramification and P2Y12 expression decrease, even as the demand for neuronal surveillance and the identification of irreversibly damaged neurons intensifies. Our study demonstrates that P2Y12 deficiency promotes an alternative form of microglia-neuron interaction. In non-infected adult P2Y12 KO mice, body-to-body interactions between neurons and microglia were more prevalent than in WT mice. Similarly, in prion-infected P2Y12 KO mice, neuronal envelopment by reactive microglia was more frequent than in prion-infected WT mice. Several lines of evidence suggest that this interaction is distinct from engulfment, the initial step of phagocytosis. First, we previously demonstrated that neuronal envelopment does not require CD11b ^33^, a mediator of phagocytosis of newborn cells during neurodevelopment ^48,49^. Second, envelopment is independent of P2Y12, which drives the engulfment of virus-infected neurons. Third, neuronal envelopment increases progressively with disease advancement but does not lead to neuronal loss ^33^. Fourth, in most instances of envelopment, microglia only partially encircle neuronal surfaces ^33^. Lastly, an increase in prevalence of neuronal engulfment in prion-infected P2Y12 KO compared to WT counterparts was not accompanied by upregulation in expression of genes linked to phagocytosis (*Trem2, Mertk, CD68, Tlr2*). We hypothesize that these close body-to-body interactions, or neuronal envelopment by reactive microglia, may represent an alternative mechanism for monitoring neuronal health or stress, especially under conditions of chronic inflammation. The prevalence of body-to-body interactions is expected to increase as neuronal dysfunction becomes more pronounced during disease progression, a pattern previously observed in prion disease ^33^.

This study is the first to implicate the P2Y12 receptor in defining an outcome of neurodegenerative disease, extending its role beyond homeostatic, ramified microglia. Because prion disease naturally affects both humans and animals, prion-inoculated mice develop authentic neurodegenerative disease state rather than modelling a disease. It is intriguing that P2Y12, often considered a homeostatic microglial marker ^9-14^, influences the very late stage of the disease. We observed P2Y12 expression at the advanced disease stages and in brain regions with intense neuroinflammation. More importantly, P2Y12 was expressed in reactive microglia with ameboid morphology, challenging the prevailing notion that its expression is linked to homeostatic microglia. While P2Y12 expression declines in individual cells, its overall expression across the microglial population increases, likely due to microglial proliferation ^30,33,35^. Notably, P2Y12 deletion shifted microglial morphology toward an ameboid shape in non-infected adult mice. Collectively, these findings suggest that P2Y12 is not only essential for process-to-body interactions but also for maintaining a balance between process-to-body and body-to-body contacts. In the broader context of neurodegeneration, the progressive decline of P2Y12 expression in individual microglia cells may serve either an adaptive or maladaptive role by lowering the threshold for forming body-to-body interactions with neurons.

While an initial increase in envelopment prevalence may serve as an adaptive response to pathological changes, our findings suggest that excessive envelopment is detrimental and accelerates disease progression. Prolonged envelopment events could disrupt neuronal functions, contributing to neurodegeneration. One of the factors leading to the prolonged envelopment could be a reduced motility of individual microglia cells resulting from a declining P2Y12 expression. In our study, isolated reactive microglia exhibited much higher basal motility than homeostatic microglia. The concurrent increase in basal motility and decline in P2Y12 expression with disease progression suggests that both P2Y12-dependent and independent mechanisms may regulate reactive microglial motility in prion-infected animals. Indeed, while motility was reduced in prion-infected P2Y12 KO mice compared to WT counterparts, it remained significantly higher than in non-infected mice. Notably, observation of high basal motility of reactive microglia in the absence of external stimuli was surprising. This observation suggests that elevated motility may be an intrinsic feature of the reactive phenotype, a possibility that warrants further investigation in future studies.

In the current work, the maximum capacity of microglia to uptake PrP^Sc^ was reached faster in P2Y12 KO mice compared to WT mice. By the time of clinical onset, more microglial cells contained PrP^Sc^ in P2Y12 KO mice than in WT counterparts. Despite this enhanced clearance of PrP^Sc^, disease progression was not mitigated but instead accelerated after clinical onset in P2Y12 KO animals. Our previous work demonstrated that by the time of clinical onset, prion replication is not controllable, as microglia sequester only a small fraction of total PrP^Sc 33^. Nevertheless, it is difficult to explain the accelerated disease progression in P2Y12 KO mice without considering the detrimental impact of excessive neuronal envelopment, which intensifies after clinical onset.

Comparison of SSLOW and 22L prion strains provides further insights. Our prior studies showed that 22L-infected mice accumulate significantly less PrP^Sc^ in microglia than SSLOW-infected mice ^33^. This difference could stem from a slower rate of 22L PrP^Sc^ uptake by microglia or faster degradation of 22L PrP^Sc^ in the lysosomal compartments relative to the uptake or degradation of SSLOW PrP^Sc^. Regardless, 22L-infected mice exhibited lower neuroinflammation and reduced neuronal envelopment compared to SSLOW-infected animals^33,41,50^. The relationship between microglial PrP^Sc^ accumulation, neuroinflammation, and neuronal envelopment remains to be clarified. It is unclear whether PrP^Sc^ uptake by microglia induces pathways responsible for neuronal envelopment. PrP^Sc^ uptake precedes neuronal envelopment, and microglial cells involved in envelopment contain PrP^Sc^ deposits ^33^. The transition to neuronal envelopment coincides with a decline in the rate of PrP^Sc^ uptake. However, whether PrP^Sc^ uptake by individual cells primes microglia for neuronal envelopment is uncertain.

Excessive accumulation of lysosomal PrP^Sc^, which resists proteolytic degradation, may contribute to lysosomal damage. Consistent with this hypothesis, the dynamics of Gal3 upregulation mirrored that of PrP^Sc^ accumulation in microglia. Gal3 has been recognized as a marker of lysosomal membrane rupture and has been implicated in lysosomal repair ^42,43^. Notably, early PrP^Sc^ accumulation and Gal3 upregulation were associated with more pronounced neuronal envelopment in P2Y12 KO mice compared to WT controls, raising a possibility of a mechanistic link between these microglial activities.

This study demonstrates that P2Y12, a key purinergic receptor, regulates microglia-neuron interactions and plays an unexpected role in prion disease. Under homeostatic conditions, P2Y12 is essential for process-to-body contacts; however, its deletion shifts microglial interactions toward body-to-body contacts. In prion-infected mice, P2Y12 deletion increases the prevalence of neuronal envelopment by reactive microglia. This excessive envelopment, likely exacerbated by reduced microglial motility following P2Y12 loss, accelerates disease progression. Notably, despite increased PrP^Sc^ uptake by microglia in P2Y12 KO mice at clinical onset, disease progression was not mitigated but rather accelerated, suggesting that extensive neuronal envelopment is detrimental. Furthermore, this study challenges the notion that P2Y12 expression is specific to ramified microglia, revealing its presence in reactive, ameboid microglia at terminal disease stages. Collectively, these findings redefine the role of P2Y12 in neurodegeneration, suggesting that its progressive decline may lower the threshold for body-to-body microglia-neuron interactions, ultimately influencing disease outcomes.

## Methods

### Animals

Brain-derived material for inoculations was prepared from terminally ill SSLOW- or 22L-infected wild type (WT, C57Bl/6J) mice as 10% (w/v) brain homogenate (BH) in PBS, pH 7.4, using glass/Teflon homogenizers attached to a cordless 12 V compact drill ^40^. Immediately before inoculation, the inoculum was diluted to 1% in PBS, pH 7.4, and further dispersed by 30 sec indirect sonication at ∼200 watts in a microplate horn of a sonicator (Qsonica, Newtown, CT). P2Y12 KO (P2Y12^-/-^) ^51,52^ and WT mice were inoculated intracerebrally with 20 μl of inoculum under 3% isoflurane anesthesia. Alternatively, C57Bl/6J mice were inoculated with 20 μl 10% BH ic (Fig. 1a, 2c) or, for Fig. 1b, c, d and S1, with 200 μl 1% BH intraperitoneally. Animals were scored weekly using four categories, each of which was graded using score ‘0 – 3’, with ‘3’ being the most severe impairment. The scoring categories were: clasping hind legs, posture (kyphosis, rigid tail, rearing difficulties), mobility (difficulties in ambulation and navigating the cage edge), and gate (keeping balance while walking, wobbly gate, disorientation, lethargy). The animals were deemed symptomatic when they displayed a consistent increase in the combined score starting from ‘4’. The mice were considered terminal and euthanized when they were unable to rear and/or lost 20% of their weight. Control groups were age-matched non-inoculated mice.

### Antibodies

Primary antibodies used for immunofluorescence, immunohistochemistry and immunoblotting were as follows: rabbit polyclonal anti-IBA1 (#013-27691, FUJIFILM Wako Chemicals USA; Richmond, VA); goat polyclonal anti-IBA1 (#NB100-1028, Novus, Centennial, CO); rabbit P2Y12 (#55043A, Anaspec, Fremont, CA); TMEM119 (#90840, Cell Signaling, Danvers, MA), mouse monoclonal anti-NeuN, clone A60 (#MAB377, Millipore Sigma, Burlington, MA); mouse monoclonal anti-prion protein, clone SAF-84 (#189775, Cayman, Ann Arbor, MI); rabbit monoclonal anti-prion protein, clone 3D17 (#ZRB1268, Millipore Sigma); rat monoclonal anti-LAMP1, clone 1D4B (#121601, BioLegend, San Diego, CA);rat monoclonal anti-Gal3, clone M3/38 (#sc-23938, Santa Cruz, Dallas, TX); rabbit anti-Vim (#5741, Cell Signaling); mouse monoclonal anti-β-actin, clone AC-15 (#A5441, Sigma-Aldrich, Saint Luis, MO); The secondary antibodies for immunofluorescence were Alexa Fluor 488-, 546-, and 647-labeled (ThermoFisher Scientific, Waltham, MA).

### DAB staining and immunofluorescence

Formalin-fixed brains (3 mm slices) were treated for 1 hour in 96% formic acid before being embedded in paraffin using standard procedures. 4 μm sections produced with Leica RM2235 microtome (Leica Biosystems, Buffalo Grove, IL) were mounted on Superfrost Plus Microscope slides (#22-037-246, Fisher Scientific, Hampton, NH) and processed for immunohistochemistry according to standard protocols. To expose epitopes, slides were subjected to 20 min of hydrated autoclaving at 121° C in Citrate Buffer, pH6.0, Antigen Retriever (#C9999, Sigma-Aldrich). For the detection of PrP^Sc^, an additional 3 min treatment in concentrated formic acid was applied.

For detection with 3,3′-diaminobenzidine (DAB), horseradish peroxidase-labeled secondary antibodies and DAB Quanto chromogen and substrate (VWR, Radnor, PA) were used.

For immunofluorescence, an Autofluorescence Eliminator Reagent (Sigma-Aldrich,) and Signal Enhancer (ThermoFisher) were used on slides according to the original protocols to reduce background fluorescence. Epifluorescence images were collected using an inverted microscope Nikon Eclipse TE2000-U (Nikon Instruments Inc, Melville, NY) equipped with an illumination system X-cite 120 (EXFO Photonics Solutions Inc., Exton, PA) and a cooled 12-bit CoolSnap HQ CCD camera (Photometrics, Tucson, AZ). Images were processed using ImageJ software (1.54f, National Institute of Health, Bethesda, MD, United States).

Full cortex images were acquired and stitched using Leica MICA widefield microscope with the 20x dry objective and a resolution of 2432 × 2032 pixels per tile. Then, the stitched images were processed with Leica’s Instant Computational Clearing (ICC). Microglia morphology analysis was done using ImageJ, the MicrogliaMorphology ImageJ tool and MicrogliaMorphologyR (R package) according to Ciernia et al ^53^, with minor adjustments. Namely, all cortex images were subjected to a default threshold of ImageJ and then additional morphology measures were acquired using ImageJ batch processing and merged with the data using RStudio. ColorByCluster, a part of the MicrogliaMorphology plugin, was used to visualize the clusters on the raw images. Two clusters were selected as the optimal number to categorize microglial cells, effectively sorting each phenotype into a distinct group.

### Quantification of envelopment and neuronal count

To estimate the percentage of neurons enveloped by microglia, brains were co-immunostained for NeuN and IBA1, and immunofluorescence images from cortices of each brain were collected under 20× objective. Using ImageJ software, NeuN channel images were subjected to a threshold and watershed, converted to binary, and then individual neurons were identified as regions of interest (ROIs) using the Analyze Particles function of ImageJ. Then, corresponding IBA1 channel images were subjected to a threshold, converted to binary, and the area of IBA1 signal in ROIs defined for individual neurons was measured. Neurons were counted as undergoing envelopment if they had over 20 pixels of NeuN signal overlapped by IBA1 signal. The total number of identified NeuN^+^ cells was recorded as a neuronal count.

To quantify neurons in the pyramidal layer of the hippocampus, NeuN immunostaining was imaged under the 20× objective, and for each image, a manual count of neurons was performed within 1 – 3 rectangles (100×200 pixels) encompassing pyramidal neurons in the CA1 region of the hippocampus.

### Confocal microscopy and 3D image reconstruction

Confocal images were acquired with Leica TCS SP8 microscope using laser lines 405, 488, 552, and 638, as needed, the 40×/1.30 or 63×/1.40 oil immersion objective, the resolution of 1024×1024 pixels, and a scan speed of 400 Hz. For 3D reconstruction, the system-optimized number of steps was used. Images were processed using the LAS X and ImageJ software.

Images of non-infected brains stained with DAPI and antibodies for IBA1 and NeuN (Fig. 3) were acquired with a scan speed of 600 Hz. 3D maximum intensity projection (MIP) images consisted of 8 stacks per image. Each MIP image was split into channels in ImageJ. IBA1 channel was converted into a binary image, and the resulting thresholded image was cross-checked with the original image to make sure that all processes were still present. Analyze Particles feature in ImageJ was used to isolate a mask of particles over 20 μm^2^, which were saved as microglial regions of interest (ROIs). Then, a median filter was applied to NeuN channel, the image was thresholded using an iterative algorithm for minimum cross entropy thresholding ^54^, and Analyze Particles feature was used to isolate neurons of over 20 μm^2^. Microglial ROIs were measured on NeuN particles to detect contacts. The number of process-to-body and body-to-body contacts in each field of view was manually counted. Shape descriptors and skeleton analysis data were acquired for microglial cells in ImageJ as morphological parameters. Principle component analysis was performed using MetaboAnalyst 6.0 software (https://new.metaboanalyst.ca/MetaboAnalyst/).

### Quantification of PrP^Sc+^ microglia

Brains were co-immunostained with anti-PrP SAF-84 and IBA1 antibodies. Confocal images were subjected to a background subtraction in ImageJ, with a despeckle step and a median filter applied as needed. IBA1 and SAF-84 channels were then thresholded using the threshold selection method from gray-level histograms ^55^ and the Intermodes method ^56^, correspondingly. A mask of microglia with an area over 20 µm^2^ was created using the Analyze Particle function and microglia were added to ROI manager. Microglial ROIs were overlaid and measured on the SAF-84 channel, and “percent area” was used to count the number of PrP^Sc^ positive and negative microglia. The number of microglia cells identified in each image was also recorded.

### Western blot

For Western blots, 10% (w/v) brain homogenates (BH) were prepared as previously described ^40^ using RIPA Lysis Buffer (Millipore Sigma, St. Louis, MO). To analyze brain-derived PrP^Sc^, BH aliquots were diluted with RIPA buffer to achieve 5% BH final concentration and treated with 20 µg/ml proteinase K (New England BioLabs) in the presence of 50 mM Tris, pH 7.5, and 2% Sarcosyl, for 30 min at 37°C. To analyze other proteins, BH was diluted with RIPA buffer to 1% and proteinase digestion was omitted. The resulting samples were supplemented with 4xSDS loading buffer and heated for 10 min in a boiling water bath before loading onto NuPAGE 12% Bis-Tris gels. Wet transfer onto PVDF membranes and probing of Western blots was done according to standard procedures. The signals were visualized by Immobilon Forte Western HRP Substrate (Millipore Sigma, Rockfield, MD) or SuperSignal West pico PLUS Chemiluminescent Substrate (Thermo Scientific, Rockford, IL) using Invitrogen iBright 1500 imager, and quantified with iBright Analysis software (Thermo Scientific, Rockford, IL). Intensity data were presented as normalized by actin, except for PrP^Sc^ Western blots treated with protease K.

### NanoString

Mice were euthanized, and their brains were immediately extracted and dissected to collect individual regions for RNA isolation using Trizol (Thermo Fisher Scientific, Waltham, MA, USA) and Aurum Total RNA Mini Kit (Bio-Rad, Hercules, CA, USA), as published ^57^. An amount of 200 ng of total RNA was submitted to the Institute for Genome Sciences at the University of Maryland, School of Medicine, for RNA integrity check and subsequent analysis using a custom nCounter Mouse Astrocyte Panel, which also included microglia-specific genes ^50,58^. Only samples with an RNA integrity number RIN > 7.2 were used for NanoString analysis. For each sample, the assessment of all target sequences was performed within a single tube, using uniquely coded hybridization probes designed by NanoString Thechnologies (Seattle, WA), enabling a reliable and reproducible assessment of the expression. All data passed quality control assessments for imaging, binding, positive control, or CodeSet content normalization.

### RT-qPCR

Total RNA was isolated from 10% BH in RIPA buffer. 100 μl BH aliquots were further homogenized within RNase-free 1.5-mL tubes in 200 μL of Trizol (Thermo Fisher Scientific, Waltham, MA, USA), using RNase-free disposable pestles (Fisher Scientific, Hampton, NH, USA). After homogenization, an additional 600 μL of Trizol was added to each homogenate, and the samples were centrifuged at 11,400× g for 5 min at 4 °C. The supernatant was collected, incubated for 5 min at room temperature, then supplemented with 160 μL of cold chloroform and vigorously shaken for 30 s by hand. After an additional 5-min incubation at room temperature, the samples were centrifuged at 11,400× g for 15 min at 4 °C. The top layer was transferred to new RNase-free tubes and mixed with an equal amount of 70% ethanol. Subsequent steps were performed using an Aurum Total RNA Mini Kit (Bio-Rad, Hercules, CA, USA) following the manufacturer instructions. Isolated total RNA was subjected to DNase I digestion. RNA purity and concentrations were estimated using a NanoDrop One Spectrophotometer (Thermo Fisher Scientific, Waltham, MA). Complementary DNA (cDNA) synthesis was performed using iScript cDNA Synthesis Kit as described elsewhere. The cDNA was amplified with CFX96 Touch Real-Time PCR Detection System (Bio-Rad, Hercules, CA) using SsoAdvanced Universal SYBR Green Supermix and the primers listed in Table S1. The PCR protocol consisted of 95 °C incubation for 2 min followed by 40 amplification cycles at 95 °C for 5 s and 60 °C for 30 s. The data were analyzed using CFX96 Touch Real-Time PCR Detection System Software.

### Live cell track analysis

Mouse microglial cells were purified using Adult Brain Dissociation Kit with CD11b antibodies according to the manufacturer protocol (Miltenyi Biotec, #130-107-677). For non-infected WT and P2Y12 KO mice, microglia were purified from pools of 3 cortices, which ensured enough cells. For animals infected with SSLOW (1% BH via i.p. route), microglia were purified from individual cortices of early clinical mice at 118 dpi, in parallel (n=3 for WT and n=2 for P2Y12 KO). Cells were plated into wells of 24-well plates and left overnight to settle. The next morning live images were taken for 6 hours with 5 min intervals. Images were taken on MICA Leica microscope in phase contrast mode and 10× magnification. During the whole imaging session, cells were kept at 37 °C in an atmosphere of 5% CO_2_. For microglia motility analysis, TrackMate plugin for Fiji ImageJ software was used. All individual tracks were manually validated for the accuracy of automated tracking and corrected if needed. Individual cell Mean Track Speed and Total Track Distance parameters were used to plot the graphs.

### Study approval

The study was carried out in strict accordance with the recommendations in the Guide for the Care and Use of Laboratory Animals of the National Institutes of Health. The animal protocol was approved by the Institutional Animal Care and Use Committee of the University of Maryland, Baltimore (Assurance

Number: A32000-01; Permit Number: 0215002). Human brain tissues have been collected under the Italian National Surveillance program for CJD and related disorders, and their use for research was approved by written informed consent of patients during life or their next of kin after death.

### Statistical analysis

Statistical analyses and plotting of the data were performed using GraphPad Prism software, versions 8.4.2 10.4.1 for Windows (GraphPad Software, San Diego, California USA) or Excel 2016 - version 2302. Statistical comparison of two groups with normal distribution of data points and equal variances was done with unpaired two-tailed t-test. Alternatively, non-parametric two-tailed Mann-Whitney test was used. In case of unequal variances, the comparison of two groups with normal distribution of data points was performed with unpaired two-tailed t-test with Welch’s correction. For statistical comparison of more than two groups, one-way ANOVAs were followed by multiple comparison tests as mentioned in Figure Legends, and the resulting P-values were reported. Heat maps were made using MetaboAnalyst 6.0 software (https://new.metaboanalyst.ca/MetaboAnalyst/).

### Data availability

All the data supporting the findings of this study are available within the paper and its supplementary information les.

## Acknowledgments

Financial support for this study was provided by National Institute of Health Grants R01 NS045585 and R01 NS129502 to IVB.

## Authors Contribution

N.M. and I.V.B. conceived and designed the study; N.M., T.S., O.B., O.M., and N.P.P., performed experiments; K.M. managed mouse colony, performed animal procedures and scored the disease signs; N.M., T.S., and O.B. analyzed the data; U.B.E., provided animal model; I.V.B. wrote the manuscript. All authors read, edited and approved the final manuscript.

## References

1 Cserép, C. et al. Microglia monitor and protect neuronal function through specialized somatic purinergic junctions. Science 367, 528–537, doi:10.1126/science.aax6752 (2020).

2 Cserép, C. et al. Microglial control of neuronal development via somatic purinergic junctions. Cell Rep 40, 111369, doi:10.1016/j.celrep.2022.111369 (2022).

3 Haynes, S. E. et al. The P2Y12 receptor regulates microglial activation by extracellular nucleotides. Nature Neuroscience 9, 1512–1519, doi:10.1038/nn1805 (2006).

4 Maeda, M., Tsuda, M., Tozaki-Saitoh, H., Inoue, K. & Kiyama, H. Nerve injury-activated microglia engulf myelinated axons in a P2Y12 signaling-dependent manner in the dorsal horn. Glia 58, 1838–1846, doi:10.1002/glia.21053 (2010).

5 Fekete, R. et al. Microglia control the spread of neurotropic virus infection via P2Y12 signalling and recruit monocytes through P2Y12-independent mechanisms. Acta Neuropathol 136, 461–482, doi:10.1007/s00401-018-1885-0 (2018).

6 Ohsawa, K. et al. Involvement of P2×4 and P2Y12 receptors in ATP-induced microglial chemotaxis. Glia 55, 604–616, doi:10.1002/glia.20489 (2007).

7 Butovsky, O. et al. Identification of a unique TGF-β–dependent molecular and functional signature in microglia. Nature Neuroscience 17, 131, doi:10.1038/nn.3599 (2013).

8 Mildner, A., Huang, H., Radke, J., Stenzel, W. & Priller, J. P2Y(12) receptor is expressed on human microglia under physiological conditions throughout development and is sensitive to neuroinflammatory diseases. Glia 65, 375–387, doi:10.1002/glia.23097 (2017).

9 Butovsky, O. & Weiner, H. L. Microglial signatures and their role in health and disease. Nature Reviews Neuroscience 19, 622–635, doi:10.1038/s41583-018-0057-5 (2018).

10 Krasemann, S. et al. The TREM2-APOE Pathway Drives the Transcriptional Phenotype of Dysfunctional Microglia in Neurodegenerative Diseases. Immunity 47, 566–581.e569, doi:10.1016/j.immuni.2017.08.008 (2017).

11 Zrzavy, T. et al. Loss of ‘homeostatic’ microglia and patterns of their activation in active multiple sclerosis. Brain 140, 1900–1913, doi:10.1093/brain/awx113 (2017).

12 Kenkhuis, B. et al. Co-expression patterns of microglia markers Iba1, TMEM119 and P2RY12 in Alzheimer’s disease. Neurobiol Dis 167, 105684, doi:10.1016/j.nbd.2022.105684 (2022).

13 Holtman, I. R. et al. Induction of a common microglia gene expression signature by aging and neurodegenerative conditions: a co-expression meta-analysis. Acta Neuropathologica Communications 3, 31, doi:10.1186/s40478-015-0203-5 (2015).

14 Maeda, J. et al. Distinct microglial response against Alzheimer’s amyloid and tau pathologies characterized by P2Y12 receptor. Brain Communications 3, doi:10.1093/braincomms/fcab011 (2021).

15 Slota, J. A., Sajesh, B. V., Frost, K. F., Medina, S. J. & Booth, S. A. Dysregulation of neuroprotective astrocytes, a spectrum of microglial activation states, and altered hippocampal neurogenesis are revealed by single-cell RNA sequencing in prion disease. Acta Neuropathologica Communications 10, 161, doi:10.1186/s40478-022-01450-4 (2022).

16 Prusiner, S. B. Prions. Proc Natl Acad Sci U S A 95, 13363–13383 (1998).

17 Legname, G. et al. Synthetic mammalian prions. Science 305, 673–676 (2004).

18 Brandner, S. et al. Normal host prion protein necessary for scrapie-induced neurotoxicity. Nature 379, 339–343 (1996).

19 Sandberg, M. K. et al. Prion neuropathology follows the accumulation of alternate prion protein isoforms after infective titre has peaked. Nat Commun 5, e4347, doi:10.1038/ncomms5347 (2014).

20 Mallucci, G. et al. Depleting Neuronal PrP in Prion Infection Prevents Disease and Reverses Spongiosis. Science 302, 871–874 (2003).

21 Rambold, A. S. et al. Stress-protective signalling of prion protein is corrupted by scrapie prions. The EMBO Journal 27, 1974–1984, doi:10.1038/emboj.2008.122 (2008).

22 Fang, C., Imberdis, T., Garza, M. C., Wille, H. & Harris, D. A. A Neuronal Culture System to Detect Prion Synaptotoxicity. PLoS Pathog 12, e1005623 (2016).

23 Fang, C. et al. Prions activate a p38 MAPK synaptotoxic signaling pathway. PLOS Pathogens 14, e1007283, doi:10.1371/journal.ppat.1007283 (2018).

24 Le, N. T. T., Wu, B. & Harris, D. A. Prion neurotoxicity. Brain Pathol 29, 263–277, doi:10.1111/bpa.12694 (2019).

25 Watts, J. C. & Prusiner, S. B. Mouse Models for Studying the Formation and Propagation of Prions. Journal of Biological Chemistry 289, 19841–19849, doi:10.1074/jbc.R114.550707 (2014).

26 Cunningham, C., Wilcockson, D. C., Campion, S., Lunnon, K. & Perry, V. H. Central and systemic endotoxin challenges exacerbate the local inflammatory response and increase neuronal death during chronic neurodegeneration. J Neurosci 25, 9275–9284, doi:10.1523/jneurosci.2614-05.2005 (2005).

27 Cunningham, C. et al. Systemic inflammation induces acute behavioral and cognitive changes and accelerates neurodegenerative disease. Biol Psychiatry 65, 304–312, doi:10.1016/j.biopsych.2008.07.024 (2009).

28 Lunnon, K. et al. Systemic inflammation modulates Fc receptor expression on microglia during chronic neurodegeneration. J Immunol 186, 7215–7224, doi:10.4049/jimmunol.0903833 (2011).

29 Kranich, J. et al. Engulfment of cerebral apoptotic bodies controls the course of prion disease in a mouse strain-dependent manner. J Exp Med 207, 2271–2281 (2010).

30 Gomez-Nicola, D., Fransen, N. L., Suzzi, S. & Perry, V. H. Regulation of microglial proliferation during chronic neurodegeneration. J Neurosci 33, 2481–2493 (2013).

31 Nakagaki, T. et al. Administration of FK506 from Late Stage of Disease Prolongs Survival of Human Prion-Inoculated Mice. Neurotherapeutics 17, 1850–1860, doi:10.1007/s13311-020-00870-1 (2020).

32 Nazmi, A. et al. Chronic neurodegeneration induces type I interferon synthesis via STING, shaping microglial phenotype and accelerating disease progression. Glia 67, 1254–1276, doi:10.1002/glia.23592 (2019).

33 Makarava, N. et al. Reactive microglia partially envelop viable neurons in prion diseases. J Clin Invest 134, doi:10.1172/jci181169 (2024).

34 Lawson, V. A. Supportive care or exhausted neglect: the role of microglia at the end stage of prion disease. J Clin Invest 134, doi:10.1172/jci186940 (2024).

35 Gómez-Nicola, D., Schetters, S. T. & Perry, V. H. Differential role of CCR2 in the dynamics of microglia and perivascular macrophages during prion disease. Glia 62, 1041–1052, doi:10.1002/glia.22660 (2014).

36 Satoh, J. et al. TMEM119 marks a subset of microglia in the human brain. Neuropathology 36, 39–49, doi:10.1111/neup.12235 (2016).

37 Zhou, Y. et al. Human and mouse single-nucleus transcriptomics reveal TREM2-dependent and TREM2-independent cellular responses in Alzheimer’s disease. Nature Medicine 26, 131–142, doi:10.1038/s41591-019-0695-9 (2020).

38 Uweru, O. J. et al. A P2RY12 deficiency results in sex-specific cellular perturbations and sexually dimorphic behavioral anomalies. J Neuroinflammation 21, 95, doi:10.1186/s12974-024-03079-7 (2024).

39 Sipe, G. O. et al. Microglial P2Y12 is necessary for synaptic plasticity in mouse visual cortex. Nature Communications 7, 10905, doi:10.1038/ncomms10905 (2016).

40 Makarava, N. et al. Stabilization of a prion strain of synthetic origin requires multiple serial passages. J Biol Chem 287, 30205–30214 (2012).

41 Makarava, N., Chang, J. C.-Y., Molesworth, K. & Baskakov, I. V. Posttranslational modifications define course of prion strain adaptation and disease phenotype. The Journal of Clinical Investigation 130, 4382–4395, doi:10.1172/JCI138677 (2020).

42 Jia, J. et al. Galectin-3 Coordinates a Cellular System for Lysosomal Repair and Removal. Dev Cell 52, 69–87.e68, doi:10.1016/j.devcel.2019.10.025 (2020).

43 Aits, S. et al. Sensitive detection of lysosomal membrane permeabilization by lysosomal galectin puncta assay. Autophagy 11, 1408–1424, doi:10.1080/15548627.2015.1063871 (2015).

44 Carroll, J. A. & Chesebro, B. Neuroinflammation, Microglia, and Cell-Association during Prion Disease. Viruses 11, doi:10.3390/v11010065 (2019).

45 Aguzzi, A. & Zhu, C. Microglia in prion diseases. J Clin Invest 127, 3230–3239, doi:10.1172/jci90605 (2017).

46 Blume, Z. I., Lambert, J. M., Lovel, A. G. & Mitchell, D. M. Microglia in the developing retina couple phagocytosis with the progression of apoptosis via P2RY12 signaling. Dev Dyn 249, 723–740, doi:10.1002/dvdy.163 (2020).

47 Wu, W. et al. In vivo imaging in mouse spinal cord reveals that microglia prevent degeneration of injured axons. Nature Communications 15, 8837, doi:10.1038/s41467-024-53218-0 (2024).

48 VanRyzin, J. W. et al. Microglial Phagocytosis of Newborn Cells Is Induced by Endocannabinoids and Sculpts Sex Differences in Juvenile Rat Social Play. Neuron 102, 435–449.e436, doi:10.1016/j.neuron.2019.02.006 (2019).

49 Wakselman, S. et al. Developmental Neuronal Death in Hippocampus Requires the Microglial CD11b Integrin and DAP12 Immunoreceptor. The Journal of Neuroscience 28, 8138, doi:10.1523/JNEUROSCI.1006-08.2008 (2008).

50 Makarava, N., Mychko, O., Chang, J. C.-Y., Molesworth, K. & Baskakov, I. V. The degree of astrocyte activation is predictive of the incubation time to prion disease. Acta Neuropathologica Communications 9, 87, doi:10.1186/s40478-021-01192-9 (2021).

51 Andre, P. et al. P2Y12 regulates platelet adhesion/activation, thrombus growth, and thrombus stability in injured arteries. J Clin Invest 112, 398–406, doi:10.1172/jci17864 (2003).

52 Bisht, K. et al. Capillary-associated microglia regulate vascular structure and function through PANX1-P2RY12 coupling in mice. Nat Commun 12, 5289, doi:10.1038/s41467-021-25590-8 (2021).

53 Kim, J., Pavlidis, P. & Ciernia, A. V. Development of a High-Throughput Pipeline to Characterize Microglia Morphological States at a Single-Cell Resolution. eNeuro 11, doi:10.1523/eneuro.0014-24.2024 (2024).

54 Li, C. H. & Tam, P. K. S. An iterative algorithm for minimum cross entropy thresholding. Pattern Recognition Letters 19, 771–776, doi:10.1016/S0167-8655(98)00057-9 (1998).

55 Otsu, N. A Threshold Selection Method from Gray-Level Histograms. IEEE Transactions on Systems, Man, and Cybernetics 9, 62–66 (1979).

56 Prewitt, J. M. & Mendelsohn, M. L. The analysis of cell images. Ann N Y Acad Sci 128, 1035–1053, doi:10.1111/j.1749-6632.1965.tb11715.x (1966).

57 Makarava, N., Chang, J. C.-Y., Molesworth, K. & Baskakov, I. V. Region-specific glial homeostatic signature in prion diseases is replaced by a uniform neuroinflammation signature, common for brain regions and prion strains with different cell tropism. Neurobiology of Disease 137, e104783, doi:10.1016/j.nbd.2020.104783 (2020).

58 Makarava, N. et al. Region-Specific Homeostatic Identity of Astrocytes Is Essential for Defining Their Response to Pathological Insults. Cells 12, doi:10.3390/cells12172172 (2023).

