## Supplementary Figures for "Knockout of P2Y12 receptor facilitates microglia-neuron body-to-body interactions and accelerates prion disease"

**Supplementary data**

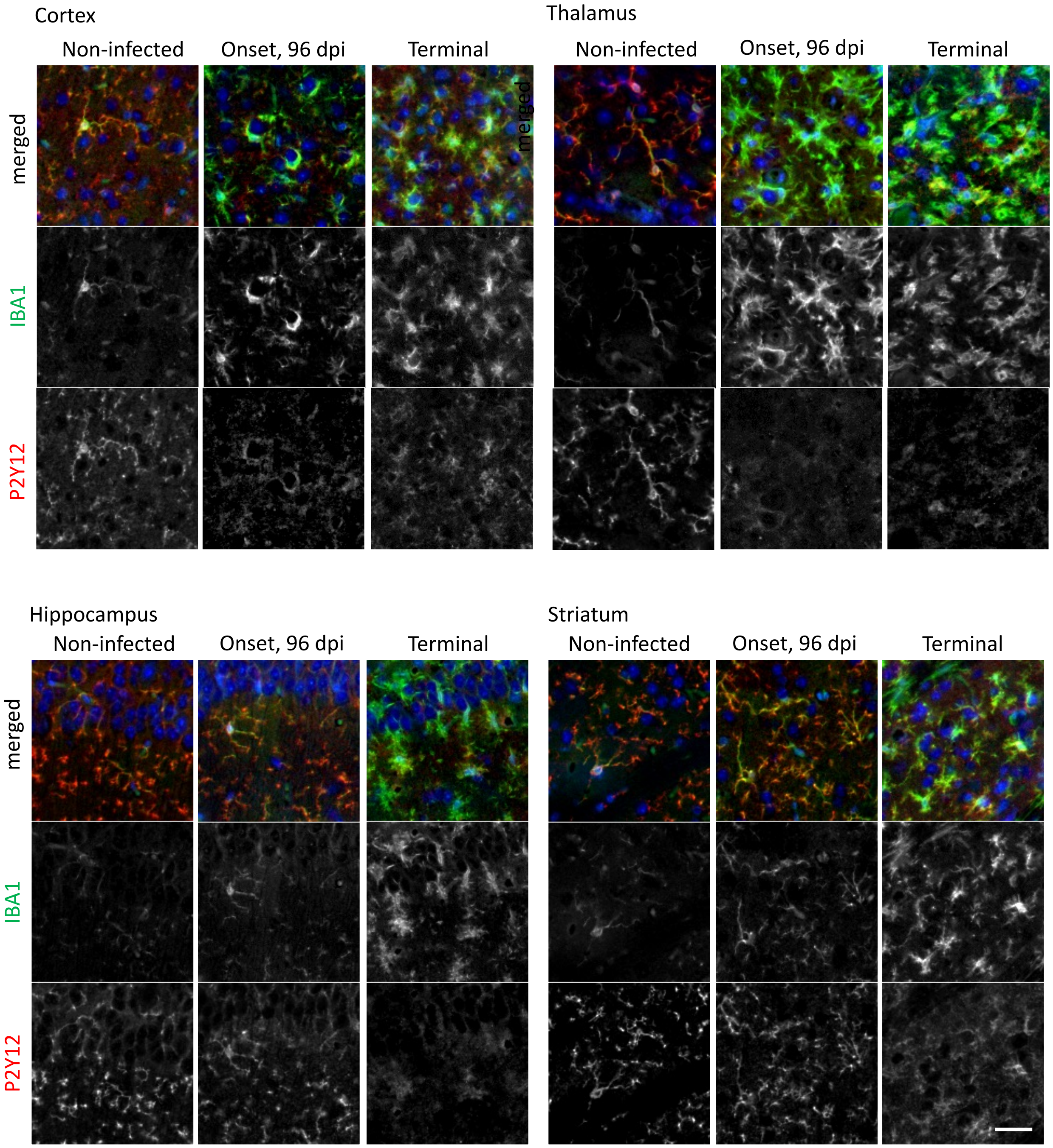


**Supplementary Fig. 1.** Magnified images of cortex, thalamus, hippocampus and striatum from non-infected WT mice and WT mice infected with 22L or SSLOW prion strains via i.p. route and examined at the terminal stage using co-immunostaining for IBA1 (upper panel) or P2Y12 (lower panel). Scale bar 20μm.

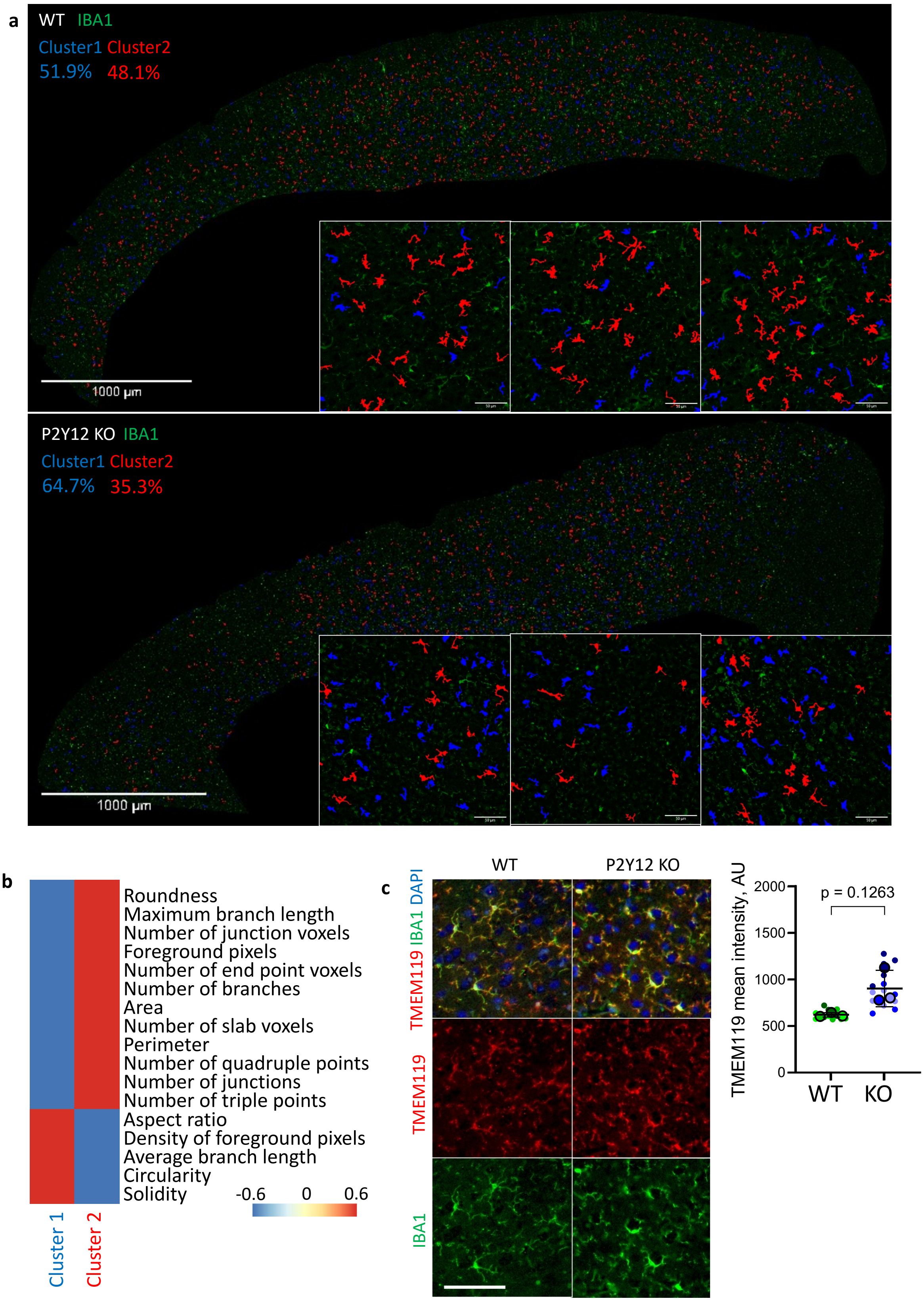
a
WT IBA1
Cluster1 Cluster2
51.9% 48.1%
1000 μm
P2Y12 KO IBA1
Cluster1 Cluster2
64.7% 35.3%
1000 μm
b
Roundness
Maximum branch length
Number of junction voxels
Foreground pixels
Number of end point voxels
Number of branches
Area
Number of slab voxels
Perimeter
Number of quadruple points
Number of junctions
Number of triple points
Aspect ratio
Density of foreground pixels
Average branch length
Circularity
Solidity
-0.6 0 0.6
Cluster 1
Cluster 2
c
WT
P2Y12 KO
TMEM119 IBA1 DAPI
TMEM119
IBA1
TMEM119 mean intensity, AU
2000
1500
1000
500
0
p = 0.1263
WT
KO

**Supplementary Fig. 2. P2Y12 deletion alters the morphology of homeostatic microglia. a.** Full-cortex scans of IBA1-immunostained WT (upper panel) and P2Y12 KO (lower panel) brains showing that microglia could be divided into two clusters according to their morphology (blue and red). Average relative abundance of each cluster shown in corresponding colors, n=3 brains per group. Insets show magnified representative views. Scale bar 1000 µm. **b.** Heat map showing a difference in morphological parameters between two clusters of microglia. **c.**TMEM119 (red) and IBA1 (green) co-immunostaining of microglia in cortex of WT and P2Y12 KO non-infected mice (left) and a comparison of TMEM119 intensities by unpaired t-test with Welch's correction, n=3 brains per group (right). Means ± SD are marked by black lines. Scale bar 50 µm.

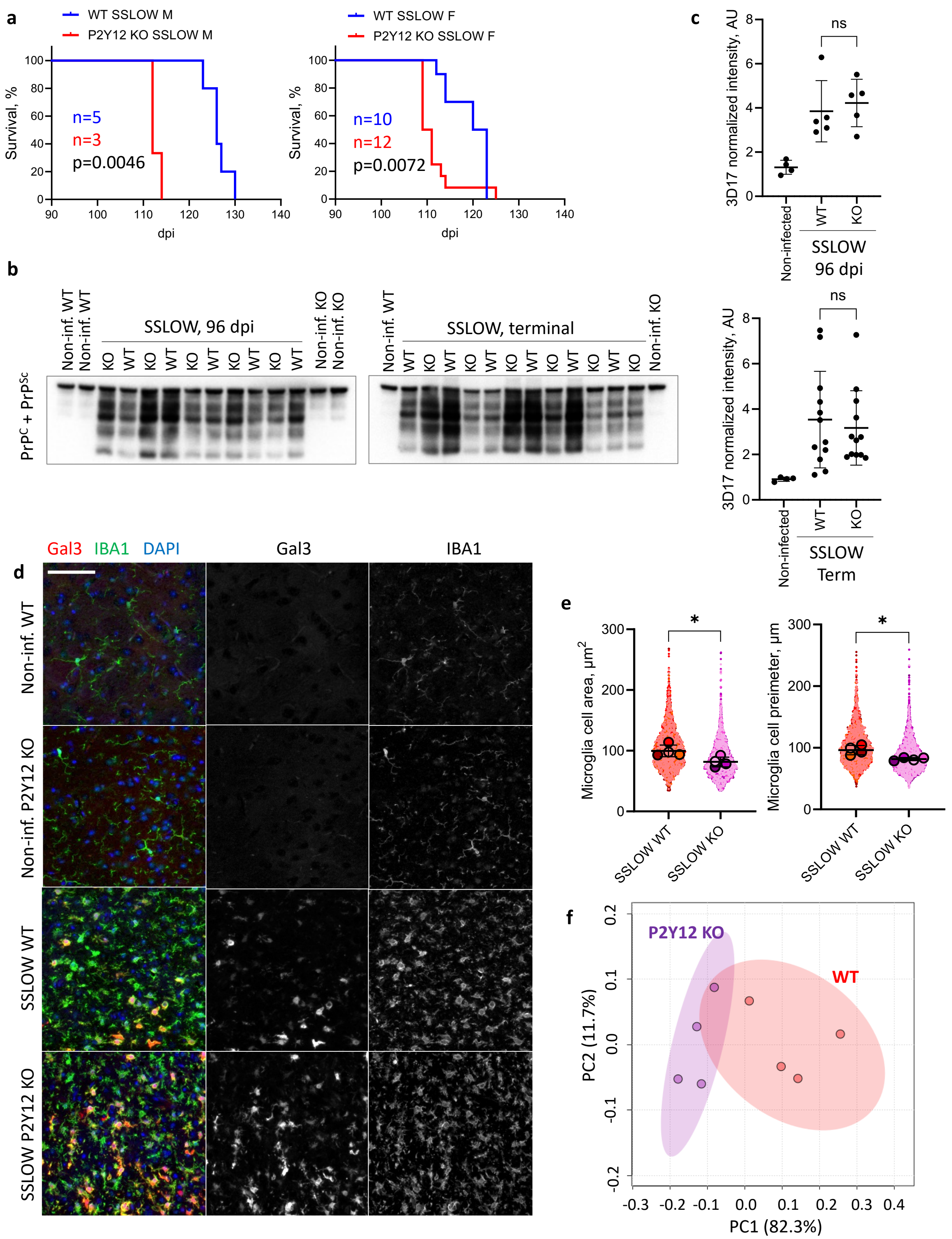

a
WT SSLOW M
P2Y12 KO SSLOW M
Survival, %
n=5
n=3
p=0.0046
dpi
WT SSLOW F
P2Y12 KO SSLOW F
n=10
n=12
p=0.0072
b
PrP$^C$ + PrP$^{Sc}$
Non-inf. WT
SSLOW, 96 dpi
Non-inf. KO
SSLOW, terminal
Non-inf. KO
c
3D17 normalized intensity, AU
ns
Non-infected
WT
KO
SSLOW
96 dpi
SSLOW
Term
Gal3 IBA1 DAPI
Gal3
IBA1
d
Non-inf. WT
Non-inf. P2Y12 KO
SSLOW WT
SSLOW P2Y12 KO
e
Microglia cell area, µm$^2$
Microglia cell preimeter, µm
SSLOW WT
SSLOW KO
f
P2Y12 KO
WT
PC2 (11.7%)
PC1 (82.3%)

**Supplementary Fig. 3. Comparison of prion-infected P2Y12 KO and WT mice. a.** Survival curves for SSLOW-infected P2Y12 KO and WT males (right) and females (left) mice. Comparison by Mantel-Cox test, n=3-12 for each group. **b.** Representative Western blot images with anti-PrP 3D17 antibody showing total PrP ($PrP^{C}$ + $PrP^{Sc}$) in brains of SSLOW-infected P2Y12 KO and WT mice at 96 dpi (left) and terminal stage of the disease (right). PrP in non-infected brains is shown as a references. For assessing total PrP, protease K digestion of sampled was omitted. **c.** Quantification of total PrP in brains of SSLOW-infected P2Y12 KO and WT mice at the onset (96 dpi, top) and terminal stage of the disease (bottom) presented as Means ± SD. SSLOW-infected P2Y12 KO and WT groups were compared by unpaired Student's t-test, n=5 per group at the onset, and n=12 per group at the terminal stage, ns – non-significant. Data for non-infected brains are shown as a reference. **d.** Confocal images of Gal3 (red) and IBA1 (green) co-immunofluorescence staining for thalami of SSLOW-infected WT and P2Y12 KO mice at the terminal stage of the disease. Scale bar 50 µm. **e.** Comparison of microglia morphology in cortices of SSLOW-infected P2Y12 KO and WT at the terminal stage by selected morphological parameters. Colors represent different brains; dots represent individual cells. Average values for each brain are shown as circles. Means ± SD are marked by black lines. Comparison by unpaired Student's t-test, n=4 brains for each group, 413 – 1695 cells per brain. *$p<0.05$. **f**. Principal component analysis of morphological parameters cortical microglia from SSLOW-infected WT and P2Y12 KO mice. Circles represent individual animals.

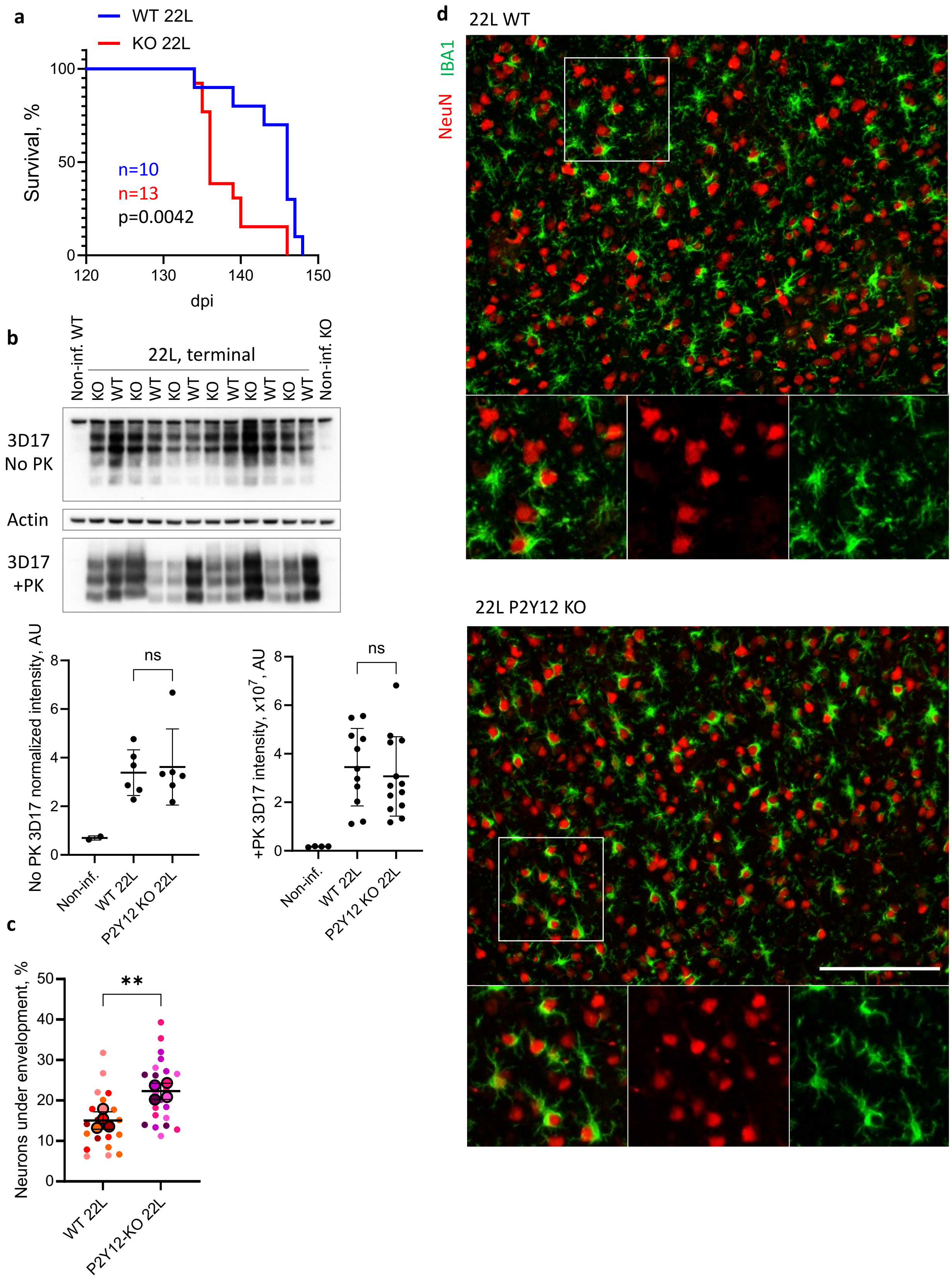
a
WT 22L
KO 22L
Survival, %
100
50
0
n=10
n=13
p=0.0042
120
130
140
150
dpi
b
Non-inf. WT
22L, terminal
Non-inf. KO
KO WT KO WT KO WT KO WT KO WT KO WT
3D17
No PK
Actin
3D17
+PK
No PK 3D17 normalized intensity, AU
ns
Non-inf.
WT 22L
P2Y12 KO 22L
+PK 3D17 intensity, x10^7, AU
ns
Non-inf.
WT 22L
P2Y12 KO 22L
c
Neurons under envelopment, %
**
WT 22L
P2Y12-KO 22L
d
22L WT
IBA1
NeuN
22L P2Y12 KO

**Supplementary Fig. 4. The effects of P2Y12 deletion in mice infected with 22L prion strain are similar to those of SSLOW-infected mice. a.** Survival curve of WT and P2Y12 KO mice infected with 22L prion strain via i.c. route. Comparison by Mantel-Cox test, n=10-13 mice (F+M) per group. **b.** Representative Western blots for total PrP (top, no proteinase K digestion) and $PrP^{Sc}$ (bottom, after proteinase K digestion) in brains of 22L-infected WT and P2Y12 KO mice at terminal stage of the disease. **c**. Quantification of the amount of total PrP (left) and $PrP^{Sc}$ (right). ns – non-significant by unpaired Student's t-test, n=6-13 mice for each group. Data for non-infected brains are shown as a reference. **d.** Quantification of neuronal envelopment in the cortices of 22L-infected WT and P2Y12 KO mice at the terminal stage. Colors represent different brains; dots represent individual fields of view; average values for each brain are shown as circles. Means ± SD are marked by black lines. **$p<0.01$ by unpaired Student's t-test, n=4 brains for each group with 4-7 fields of view for each brain. Scale bar 100 µm. **e.** Immunostaining of 22L-infected brains for microglia (IBA1, green) and neurons (NeuN, red) showing envelopment in WT (top) and P2Y12 KO cortices (bottom) at terminal stage of the disease. Insets present magnified images as merged and separated channels. Scale bar 100 µm.

**Supplemental Table 1**. Primer sequences for qRT-PCR

| Primer | Accession number | Sequence |
|---|---|---|
| MERTK | NM_008587 | F 5′- ATCATCCTCGGCTGCTTCTGTG –3’ |
| | | R 5′- ACGACCAGTTGGGAATCCTCCT -3’ |
| P2RY6 | NM_183168 | F 5′- CAGTCTTTGCTGCCACAGGCAT -3’ |
| | | R 5′- AGCAAGAAGCCGATGACCGTGA -3’ |
| P2RY13 | NM_028808 | F 5′- TGGCATCAGGTGGTCAGTCACA -3’ |
| | | R 5′- TTGTGCCTGCTGTCCTTACTCC -3’ |
| P2RX7 | NM_011027 | F 5′- GAACACGGATGAGTCCTTCGTC -3’ |
| | | R 5′- CAGTGCCGAAAACCAGGATGTC -3’ |
| P2RX4 | NM_011026 | F 5′- GCTTTCAGGAGATGGCAGTGGA -3’ |
| | | R 5′- TGTAGCCAGGAGACACGTTGTG -3’ |
| Trem2 | NM_031254 | F 5′- CTACCAGTGTCAGAGTCTCCGA -3’ |
| | | R 5′- CCTCGAAACTCGATGACTCCTC -3’ |
| TMEM119 | NM_146162 | F 5′- ACTACCCATCCTCGTTCCCTGA -3’ |
| | | R 5′- TAGCAGCCAGAATGTCAGCCTG -3’ |
| TLR2 | NM_011905 | F 5′- ACAGCAAGGTCTTCCTGGTTCC -3’ |
| | | R 5′- GCTCCCTTACAGGCTGAGTTCT -3’ |
| IL1a | NM_010554 | F 5′- ACGGCTGAGTTTCAGTGAGACC -3’ |
| | | R 5′- CACTCTGGTAGGTGTAAGGTGC -3’ |
| CXCL10 | NM_021274 | F 5′- ATCATCCCTGCGAGCCTATCCT -3’ |
| | | R 5′- GACCTTTTTTGGCTAAACGCTTTC -3’ |
| CCL3 | NM_011337 | F 5′- ACTGCCTGCTGCTTCTCCTACA -3 |
| | | R 5′- ATGACACCTGGCTGGGAGCAAA -3 |
| CCL5 | NM_013653 | F 5′- CCTGCTGCTTTGCCTACCTCTC -3’ |
| | | R 5′- ACACACTTGGCGGTTCCTTCGA -3’ |
| GAPDH | NM_008084 | F 5′- CATCACTGCCACCCAGAAGACTG -3’ |
| | | R 5′- ATGCCAGTGAGCTTCCCGTTCAG -3’ |
| C3 | NM_009778 | F 5′- CGCAACGAACAGGTGGAGATCA -3’ |
| | | R 5′- CTGGAAGTAGCGATTCTTGGCG -3’ |
